# m^6^A governs length-dependent enrichment of mRNAs in stress granules

**DOI:** 10.1101/2022.02.18.480977

**Authors:** Ryan J. Ries, Brian F. Pickering, Hui Xian Poh, Sim Namkoong, Samie R. Jaffrey

**Author notes:** Correspondence should be addressed to S.R.J.

## Abstract

Stress granules are biomolecular condensates composed of protein and mRNA. Long mRNAs are enriched in stress granules, which is thought to reflect the ability of long mRNAs to form multiple RNA-RNA interactions with other mRNAs. RNA-RNA interactions are thus thought to be critical for stress granule formation. Stress granule-enriched mRNAs also often contain multiple *N*^6^-methyladenosine (m^6^A) residues. YTHDF proteins bind m^6^A, creating mRNA-protein complexes that partition into stress granules. Here we determine the basis of length-dependent enrichment of mRNAs in stress granules. We show that depletion of m^6^A is sufficient to abrogate the length-dependent enrichment of mRNAs in stress granules. We show that the presence of m^6^A predicts which mRNAs are enriched. m^6^A formation is triggered by long exons, which are often found in long mRNAs, accounting for the link between m^6^A, length and stress granule enrichment. Thus, length-dependent enrichment of mRNAs in stress granules is driven by YTHDF-mRNA interactions.

## INTRODUCTION

Stress granules are biomolecular condensates that form in response to cellular stress^1–3^. As is seen with several other intracellular condensates^4,5^, stress granules contain both protein and RNA. The RNA composition has been explored by transcriptomic analysis of stress granules, which has demonstrated that essentially all cellular mRNAs can be found to some extent in stress granules^6,7^. However, some cellular mRNAs show higher or lower enrichment in stress granules relative to their cytoplasmic concentration. These data suggest that mRNA has a general propensity to be found in stress granules but that additional unknown factor(s) can further enhance mRNA enrichment in stress granules.

Enrichment of mRNAs in stress granules during stress may serve to prevent their translation during stress. Additionally, their release after stress could be important for shaping post-stress translational responses^8^. Thus, a major goal has been to identify the features of mRNAs that are enriched in stress granules. The initial study of the stress granule transcriptome by Khong et al. suggested that mRNAs enriched in stress granules had low translational efficiency^6^. A subsequent study by Namkoong et al. found that transcripts with AU-rich elements were enriched in stress granules^7^. However, the most significant predictor of mRNA enrichment in stress granules identified in these and later studies is transcript length^6,7,9^.

The basis for the length-dependent enrichment of mRNAs in stress granules is not clear. This length effect was also seen when studying purified mRNA, which can undergo phase separation in vitro to form RNA-only condensates^9^. Since a length effect was seen with purified RNA, the length effect was attributed to “RNA self-assembly,” i.e. weak intermolecular RNA-RNA interactions, which would promote RNA condensate formation. Since longer RNAs would have more locations for intermolecular RNA-RNA interactions, longer RNAs would exhibit multiple RNA-RNA interactions and therefore preferentially associate or be retained in RNA condensates. This work has led to the idea that long RNAs might be enriched in stress granules due primarily to their ability to form multiple RNA-RNA interactions^9,10^.

Recently, the number of *N*^6^-methyladenosine (m^6^A) sites on an mRNA was also shown to correlate with greater than average mRNA enrichment in stress granules^11^. m^6^A is a modified nucleotide formed co-transcriptionally on mRNA that is most often associated with mRNA instability^12–16^. m^6^A binds the cytosolic YTHDF paralogs (DFs)^16–18^, which are enriched in stress granules^11,19^. RNAs that contain multiple m^6^A sites, rather than one m^6^A site, induce proximity and phase separation of DF proteins in vitro, suggesting that polymethylated RNAs can cause DF-mRNA complexes to preferentially partition into stress granules^11,20^. Consistent with this, DF proteins fail to efficiently localize to stress granules in m^6^A-deficient embryonic stem cells^11^. mRNA in stress granules also show higher levels of m^6^A than the total cellular mRNA pool, further suggesting that m^6^A could guide mRNAs into stress granules due to their ability to bind DF proteins^11^. Although these studies did not deplete m^6^A and determine if this leads to impaired mRNA enrichment in stress granules, the overall data supports a model in which m^6^A functions to enhance mRNA targeting to stress granules^11^.

Establishing the relative contribution of m^6^A versus transcript length in mediating mRNA enrichment in stress granules is difficult. This is because a defining feature of many m^6^A-modified transcripts is their significant length. Notably, transcript length itself does not trigger methylation—instead, methylation is associated with the presence of long exons^15,21,22^, which are often present in long transcripts. Long exons may trigger m^6^A formation since these exons are associated with unique histone modifications or RNA polymerase II modifications which may directly recruit the m^6^A writer complex^23–25^. Since m^6^A is preferentially enriched in long mRNAs, it is also possible that the high levels of m^6^A in stress granules are simply a consequence of long mRNAs being enriched in stress granules. Thus, length alone could mediate the enrichment of mRNAs in stress granules, and although m^6^A happens to be found in these mRNAs, m^6^A may have little or no significant role in mediating stress granule enrichment.

Here, we show that m^6^A shapes the transcriptome of stress granules. Using cells lacking m^6^A, we show that m^6^A directly promotes that enrichment of mRNAs into stress granules, in a manner proportional to the number of m^6^A sites. However, m^6^A is also accounts for the enrichment of long mRNAs in stress granules. Long mRNAs markedly enriched in stress granules, but this length effect is largely lost in cells lacking m^6^A. We show that the loss of long mRNA enrichment in stress granules after m^6^A depletion cannot be explained by a global reduction in transcript length in the cell. Instead, this effect is likely explained by DF proteins which bind to m^6^A-modified mRNAs and are enriched in stress granules. Thus, we propose that the explanation for the enrichment of long mRNAs in stress granules is due primarily to their increased propensity to contain large amounts of m^6^A and consequently bind DF proteins. These studies demonstrate that a function for m^6^A is to bias the composition of stress granules towards long mRNAs.

## RESULTS

### Designing a system for complete, inducible loss of m^6^A in differentiated cells

We previously showed that the DF proteins and m^6^A-containing mRNAs are enriched in stress granules in mouse embryonic stem cells (mESCs)^11^. mESCs are a useful model for studying the effects of m^6^A because nearly all other cell types are not viable upon deletion of *Mettl3* or *Mettl14*^26^, which together form the heterodimeric complex responsible for ‘writing’ m^6^A in the transcriptome. In contrast, mESCs survive even after near-complete depletion of m^6^A^22,27^. We used an mESC line that shows a complete loss of m^6^A upon deletion of *Mettl3* or *Mettl14*^27^. It should be noted that another mESC line was originally described as a *Mettl3* knockout^22^, but it is now known to express a shortened, partially functional METTL3 protein, and only shows a partial loss (∼60%) of m^6^A levels^22,26^. Our previous studies used the full *Mettl14* knockout mESC line which showed that m^6^A in mRNA is required for DF enrichment in heat shock and arsenite-induced stress granules^11,27^.

Although mESCs tolerate full knockout of *Mettl3*, they are problematic since they do not form stress granules in response to many standard stress granule-inducing stimuli^11,27^. Therefore, the role of m^6^A in mediating recruitment of DF proteins to stress granules after other types of stress is not clear. Although stable *Mettl3* deletion prevents growth of most cell types besides mESCs, we thought that induced deletion of *Mettl3* in differentiated cells may be possible. We therefore developed mouse embryonic fibroblasts (MEFs) that undergo tamoxifen-induced *Mettl3* knockout, since MEFs are commonly used to study stress granules^7,28–30^. We immortalized MEFs from *Mettl3^flox/flox^* E13.5 embryos^31^ and then infected them with a Cre-ERT2-expressing lentivirus. Clonal cell lines were isolated that express Cre in a tamoxifen-dependent manner. The resulting *Mettl3* conditional knockout cell line shows near-complete loss of METTL3 after 7 d of 4-hydroxytamoxifen (4-OHT) treatment (**Fig. 1a, Extended Data Fig. 1a**). Additionally, these cells show a near-complete loss of m^6^A in twice poly(A)-selected mRNA (**Extended Data Fig. 1b**). We also observed a large reduction in mapped reads to *Mettl3* after performing RNA-seq (**Extended Data Fig. 1c-e**). These data demonstrate that both the loss of functional METTL3 protein and a near-complete loss of m^6^A are achievable using this conditional *Mettl3* knockout cell line.

**Fig. 1:**
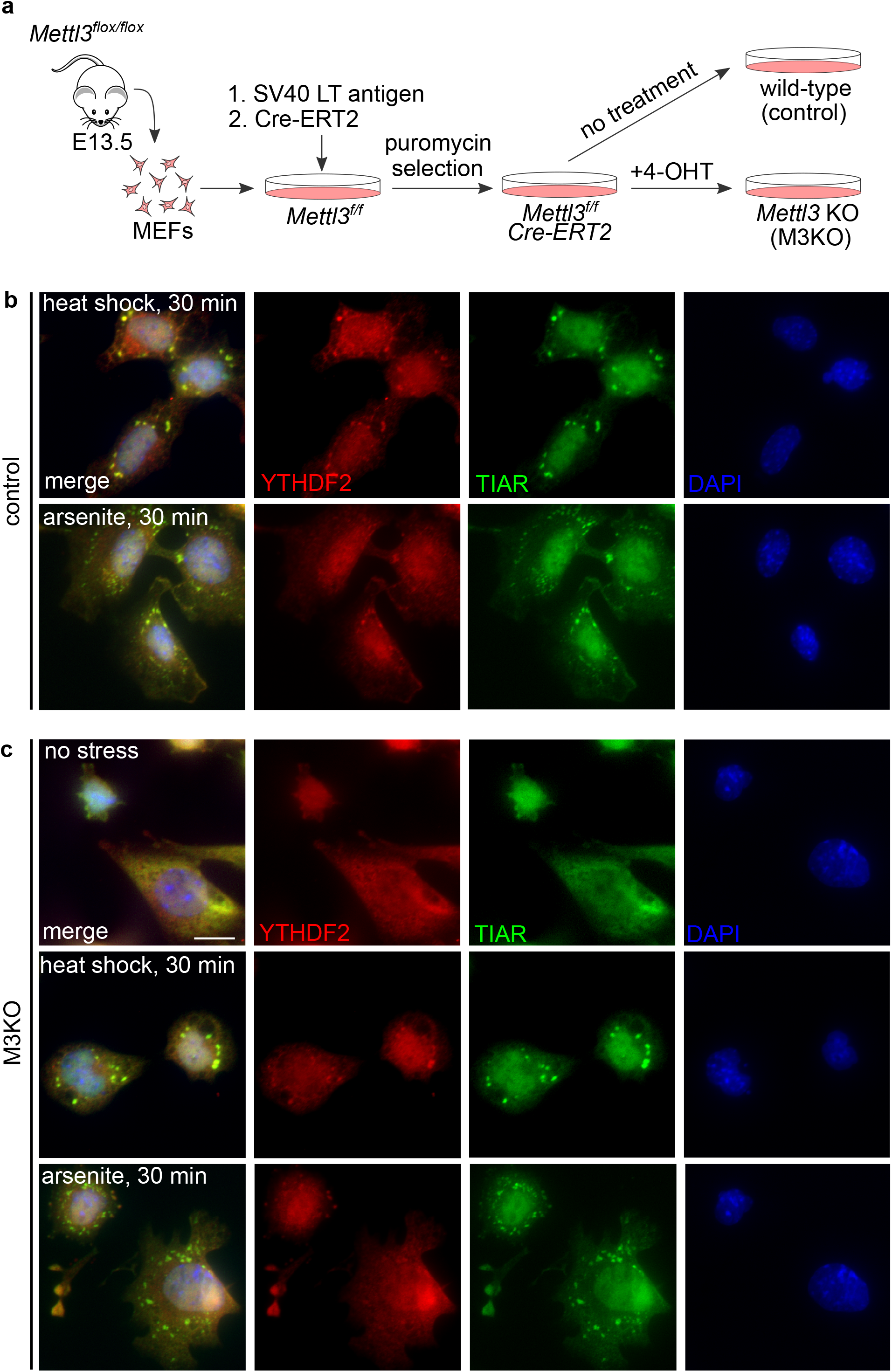
Knockout of *Mettl3* and validation of stress response in M3KO MEFs. **a,** Schematic for the design of *Mettl3* tamoxifen-inducible knockout cell line. MEFs were isolated from *Mettl3 flox/flox* mice at E13.5, immortalized with SV40 LT antigen, and transformed with *Cre-ERT2* via lentivirus. Puromycin was used to select immortalized *Cre-ERT2* positive cells. Untreated cells are considered wild-type (control) while cells treated with 4-hydroxytamoxifen (4-OHT) are considered *Mettl3* knockout (M3KO). **b,** Response to heat shock and arsenite stress in control MEFs. Heat shock was performed for 30 min at 42°C. Arsenite treatment (0.5 mM) was performed for 30 min at 37°C. YTHDF2 is depicted in red, TIAR is depicted in green, and DAPI staining is depicted in blue. **c,** Response to heat shock and arsenite stress in *Mettl3* KO cells. Conditions are as described in **b**.

### The efficiency of m^6^A-dependent DF recruitment to stress granules in MEFs is affected by the type of stress

We next wanted to determine if DF proteins are recruited to stress granules induced by different stresses in MEFs and if this effect is dependent on m^6^A. Following 7 d of either vehicle (control) or 4-OHT treatment (*Mettl3* knockout; M3KO), we applied a variety of stresses to the cells, including oxidative stress via arsenite, thermal stress via heat shock, endoplasmic reticulum stress via thapsigargin, and osmotic stress via sorbitol^29,32–34^. Stress granules were identified from immunofluorescence images by analyzing the signal of TIAR, a canonical stress granule marker^32^. In each case, we confirmed that stress granule formation occurred, even in the absence of m^6^A (**Fig. 1b,c**; **Extended Data Fig. 1f,g**). We also observed little effect of METTL3 depletion on the overall area of stress granules (**Fig. 2a,b**). These data were consistent with our previous findings in mESCs where DFs from *Mettl14* knockout mESCs showed generally normal stress granule formation after arsenite treatment and heat shock^11^.

**Fig. 2:**
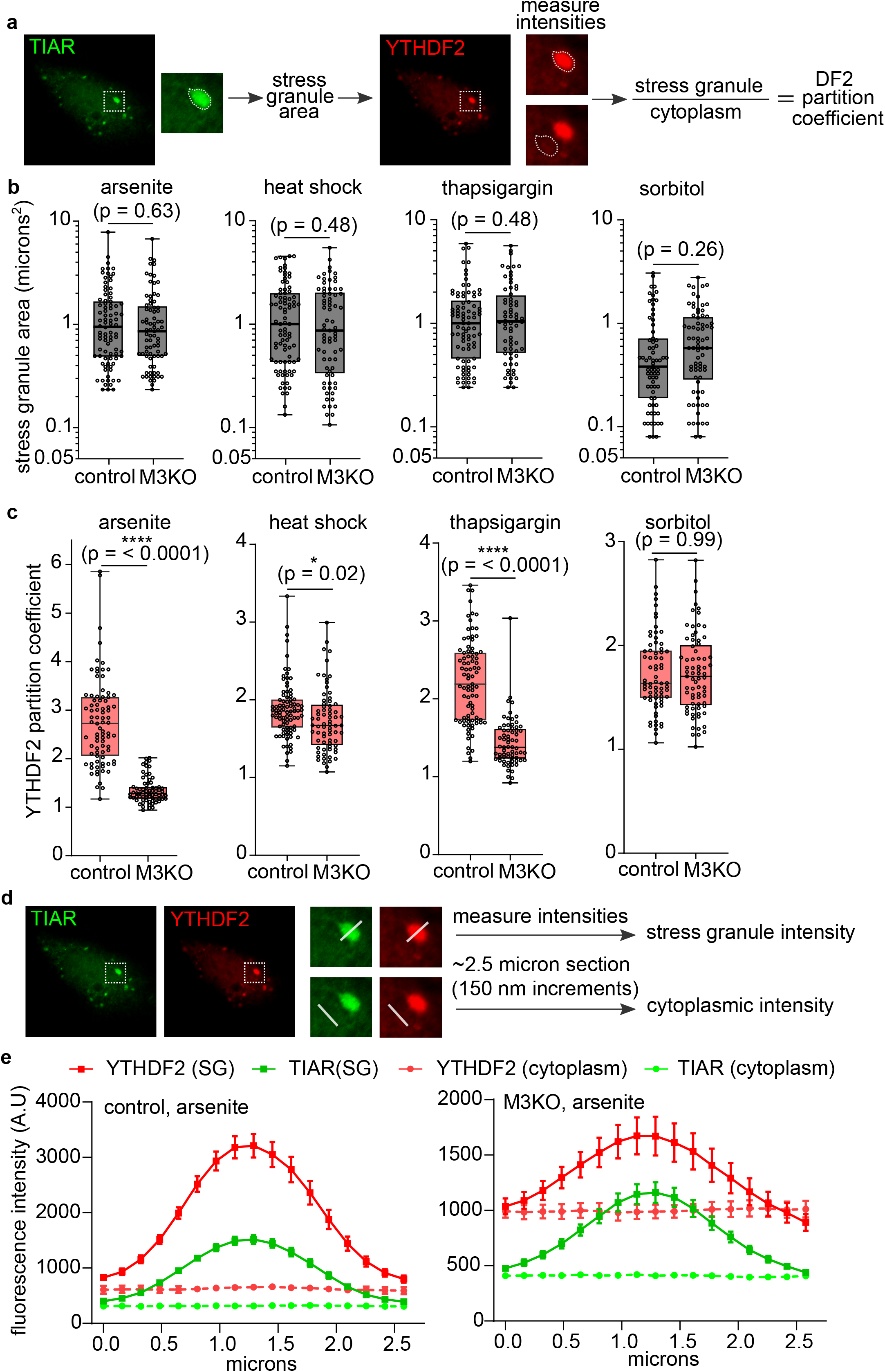
Quantification of stress differences in control and *Mettl3* knockout cells. **a,** Schematic depicting the method for measuring stress granule area and DF2 partition coefficients. TIAR fluorescence is used to define granule area which is presented in **b**. This region is then used to measure fluorescence intensities for the YTHF2 partition coefficients presented in **c**. **b,** Stress granule area measurements in control and *Mettl3* KO cells. Measurements of stress granule area show no significant differences between control and *Mettl3* KO under the stresses indicated. Boxplots depict the median and upper and lower quartiles. Open circles depict individual measurements for stress granules. Arsenite, control: n = 86, arsenite, *Mettl3* KO: n = 77; heat shock, control: n = 87; heat shock, *Mettl3* KO: n = 72; thapsigargin, control: n = 86; thapsigargin, *Mettl3* KO: n = 68; sorbitol, control: n = 71; sorbitol, *Mettl3* KO: n = 73. **c,** YTHDF2 stress granule partition coefficients in control and *Mettl3* KO. Measurements of YTHDF2 partition coefficients show no significant change with sorbitol treatment, but are significantly lowered with arsenite, thapsigargin, and thermal stress. Boxplots depict the median and upper and lower quartiles. Open circles depict individual measurements for stress granules. Arsenite, control: n = 83, arsenite, *Mettl3* KO: n = 70; heat shock, control: n = 88; heat shock, *Mettl3* KO: n = 73; thapsigargin, control: n = 87; thapsigargin, *Mettl3* KO: n = 68; sorbitol, control: n = 71; sorbitol, *Mettl3* KO: n = 73. **d,** Schematic depicting the method for collecting data depicted in **e** and Extended Data Fig 2. The two left-most panels show a representative image from which intensity data is collected. The dotted box denotes the stress granule of interest to be measured. The magnified panels on the right show the section to be measured (white line, ∼2.5 um) in the stress granule and the immediately adjacent cytoplasmic region. From the 2.5 um region of interest, 17 intensity values are recorded at each ∼150 nm increment. The resulting intensity values from both stress granules and cytoplasm are then graphed at the appropriate increments along the region of interest as shown. **e,** Partition coefficient analysis of YTHDF2 in control and *Mettl3* KO cells. The left panel shows the partition coefficient data from control cells, while the right panel shows the same data from *Mettl3* KO cells. Each point shows the mean and standard deviation for datapoints collected across several granules (control, n = 11; *Mettl3* KO, n = 13). DF2 partition coefficients are substantially higher in control relative to *Mettl3* KO stress granules. TIAR traces are shown to depict the bounds of the granule that was measured.

We were previously unable to elicit robust stress granule induction in response to thapsigargin or sorbitol treatment in mESCs^11^. This prevented an assessment of whether DF proteins are recruited to stress granules in an m^6^A-dependent manner for these stresses. Since these stresses robustly induce stress granules in MEFs, we quantified the stress granule partition coefficient for YTHDF2 (DF2), the most highly expressed m^6^A reader in most cells^35^. The partition coefficient is defined as the ratio between the fluorescence intensity in a region relative to another. The DF2 partition coefficient within stress granules was measured by determining the average pixel intensity within the bounds of the granule as defined by TIAR staining divided by the average pixel intensity of cytoplasmic DF2 in an adjacent region (**Fig. 2a**).

Using this approach, we found that m^6^A-dependent DF2 enrichment in stress granules was dependent on the stress condition. All stresses increased the DF2 partition coefficient in control and *Mettl3* knockout cells (**Fig. 2c**). For most stresses, DF2 partitioning was m^6^A dependent, since the DF2 partition coefficient was reduced in *Mettl3* knockout cells. This effect was most clear in stress granules induced by sodium arsenite (oxidative stress) but was also evident to a lesser extent in stress granules induced by heat shock and thapsigargin treatment (ER stress) (**Fig. 2c**). In contrast, stress granules induced by sorbitol treatment (osmotic stress), which is associated with a distinct protein composition from other stress granules^36,37^, showed much less DF2 enrichment relative to other stresses, and this effect showed no dependence on *Mettl3* (**Fig. 2c**). Thus, our data suggests that m^6^A promotes recruitment of DF2 proteins to most types of stress granules, with the most prominent effect seen in stress granules induced by arsenite treatment.

Another method of quantifying protein enrichment in stress granules involves measuring the fluorescence intensity of pixels across a fixed length through the stress granule (**Fig. 2d**). This method allows for the collection of multiple datapoints for a single granule by measuring intensity at fixed intervals along an axis drawn through the center of a granule. Measurement of cytoplasmic fluorescence adjacent to the granule along the same axis can then be used as a normalizing factor. Measurements of DF2 using this method corroborated the findings we made by measuring average fluorescence intensity in stress granules (**Fig. 2e**; **Extended Data Fig. 2a-c**). Again, *Mettl3* knockout cells showed a significant decrease in DF2 enrichment in stress granules with the strongest effect observed after arsenite treatment (**Fig. 2e**).

### Effects of m^6^A on recruitment of mRNAs to stress granules

Although we previously showed that m^6^A-modified mRNAs are enriched in RNAs found in stress granules^11^, it is not clear if m^6^A itself drives the enrichment of mRNAs to stress granules or if other factors, such as mRNA length drives enrichment, and m^6^A is simply present on these longer mRNAs. To test the role of m^6^A more directly, we purified arsenite-induced stress granules as described previously^7,11^, and then performed RNA-seq on mRNAs that are enriched in stress granules from both control and *Mettl3* knockout cells.

To measure how m^6^A affects the enrichment of mRNAs in stress granules, we separately analyzed mRNAs with 0, 1, 2, 3, and 4 or more annotated m^6^A sites using a previously described map of m^6^A sites in MEFs^27^. For each transcript, we first measured the amount of each mRNA in stress granules from control and *Mettl3* knockout cells. Surprisingly, we observed that mRNAs that are normally methylated showed little change in their abundance in stress granules from control compared to *Mettl3* knockout cells (**Fig. 3a**, **Extended Data Fig. 3a**). These data could indicate that m^6^A does not influence the enrichment of mRNA into stress granules.

**Fig. 3:**
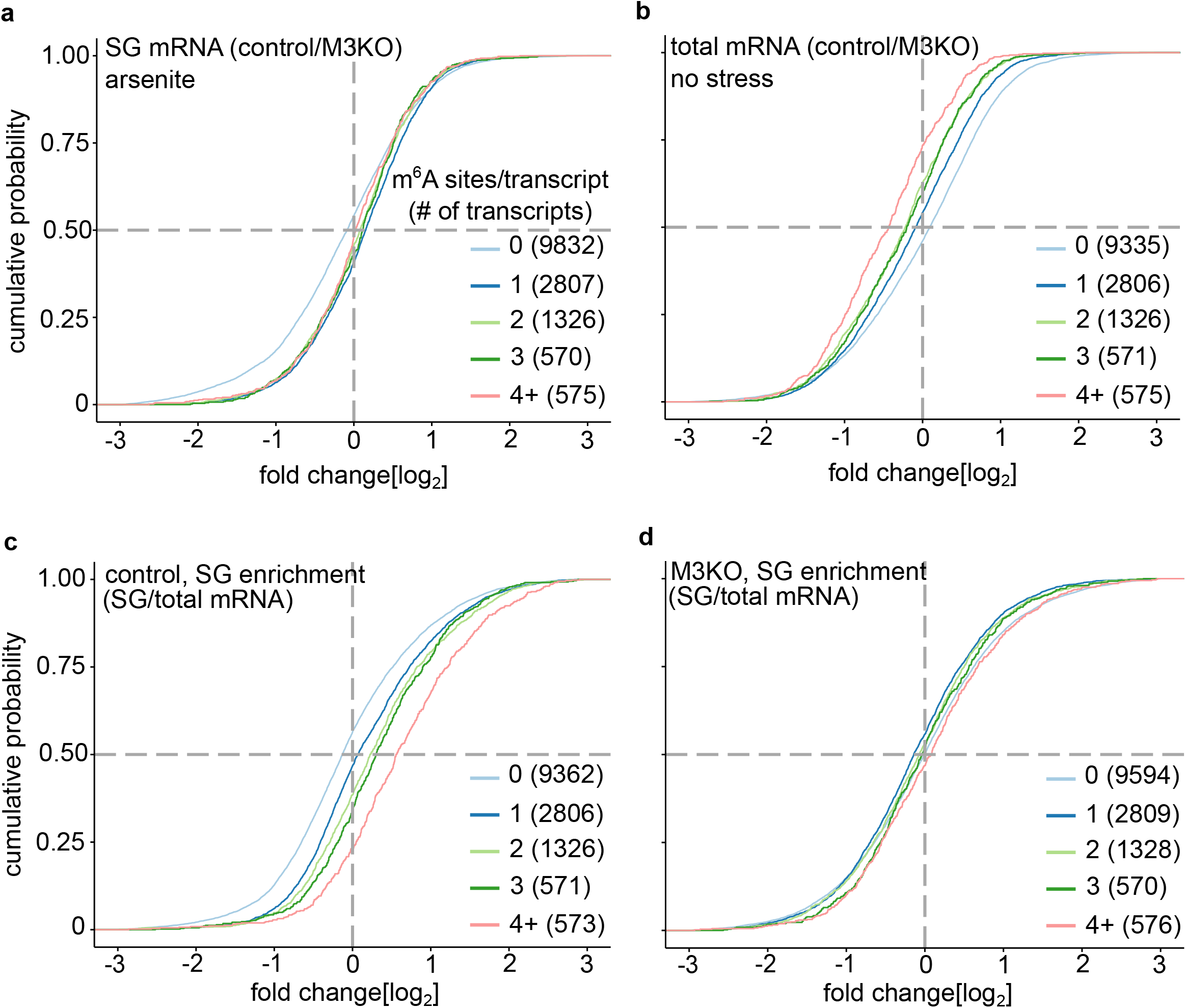
Effects of m^6^A loss on total and stress granule mRNA abundance. **a,** Cumulative distribution of log_2_ fold change for transcripts in control and *Mettl3* KO stress-granule enriched mRNAs after arsenite treatment. Control is normalized to *Mettl3* KO so that the effects of m^6^A can be visualized. m^6^A-containing transcripts are slightly enriched in control stress granules compared to *Mettl3* KO. **b,** Cumulative distribution of log_2_ fold change for transcripts in control and *Mettl3* KO total mRNA samples. The abundance of non-m^6^A containing mRNAs is relatively unchanged between control and *Mettl3* KO, while the abundance of m^6^A-containing transcripts is substantially reduced in control due to the destabilizing effects of m^6^A. **c,** Cumulative distribution of log_2_ fold change for mRNAs in stress granules compared to total mRNAs in control cells. m^6^A-containing transcripts are enriched in arsenite-induced stress granules, and the effect size increases with the number of m^6^A sites. **d,** Cumulative distribution of log_2_ fold change for mRNAs in stress granules compared to total mRNAs in *Mettl3* KO cells. The loss of m^6^A results in the loss of m^6^A-mediated mRNA enrichment in arsenite-induced stress granules.

A potential problem with this analysis is that transcripts that are normally m^6^A-modified are more highly expressed in *Mettl3* knockout cells^22,27^. This change in transcript expression is due to the loss of the degradation-inducing effect of m^6^A mRNA in knockout cells^12–16^. Indeed, m^6^A-modified mRNAs are increased in *Mettl3* knockout cells compared to control and the destabilizing effect of m^6^A is proportional to the number of m^6^A sites in the transcript (**Fig. 3b**, **Extended Data Fig. 3b**). Therefore, the high expression level of normally-methylated mRNAs in *Mettl3* knockout cells must be considered when examining the role of m^6^A in mRNA enrichment in stress granules.

We therefore reanalyzed the mRNA levels in stress granules by accounting for the markedly increased overall cellular expression of normally methylated mRNAs in *Mettl3* knockout cells. To do this, we measured an mRNA enrichment value for each mRNA by calculating mRNA abundances in stress granules relative to the total amount of each transcript in the cell. Here we observed that methylated mRNAs show a clear enrichment in control stress granules, as expected (**Fig. 3c**, **Extended Data Fig. 3c**). This is consistent with data from previous stress granule transcriptome studies which also show an m^6^A-dependent effect on mRNA enrichment in stress granules^6,7^ (**Extended Data Fig. 3e,f**). Additionally, a previous analysis showed there is a strong correlation (r = ∼0.7-0.9) between relative transcript abundances in stress granules after arsenite, heat shock, and thapsigargin stress in NIH3T3 cells, a MEF-derived cell line^7^. Thus, the mRNA recruitment mechanisms associated with arsenite-induced stress granules likely apply to mRNA recruitment to stress granules induced by these other stresses, consistent with other reports^36,37^. However, there was a striking lack between the number of m^6^A sites and mRNA enrichment in stress granules from Mettl3 knockout cells (**Fig. 3d**, **Extended Data Fig. 3d**). Thus, when considering the increased abundance of normally-methylated mRNAs in *Mettl3* knockout cells, these mRNAs no longer show preferential enrichment in stress granules. Overall, these data suggest the enrichment of m^6^A-containing mRNAs in stress granules is dependent on m^6^A.

### m^6^A mediates the length-dependent enrichment of mRNAs in stress granules

Although we found that m^6^A promotes enrichment of mRNAs in stress granules, another important determinant is transcript length^6,7,9^. Longer mRNAs are disproportionally enriched in stress granules, and genes that express transcripts with alternate 3’ UTR isoforms show preferential enrichment for the longer isoform in stress granules^7^. The ability of long mRNAs to enrich in stress granules is thought to be due to their ability to form weak and transient RNA-RNA interactions through short complementary stretches of nucleotides between two mRNAs^9,10^. This is supported by experiments that show that mRNAs can phase separate in vitro in the absence of proteins, with longer mRNAs showing greater enrichment in these protein-free RNA condensates^9^. Thus, RNA length can confer specific physical properties that could explain the enrichment of long RNAs in stress granules^10,38^.

However, it is possible that longer mRNAs are enriched in stress granules because they contain disproportionate amounts of m^6^A. Binning mRNAs by transcript length shows that the number of m^6^A sites in a transcript has a positive correlation with transcript length (**Fig. 4a**). The preferential addition of m^6^A to long mRNAs may relate to the ability of the m^6^A writer complex to be recruited to long exons^15,21,22^. Since high m^6^A levels and increased mRNA length correlate, it is difficult to disentangle the effects of these closely related transcript features.

**Fig. 4:**
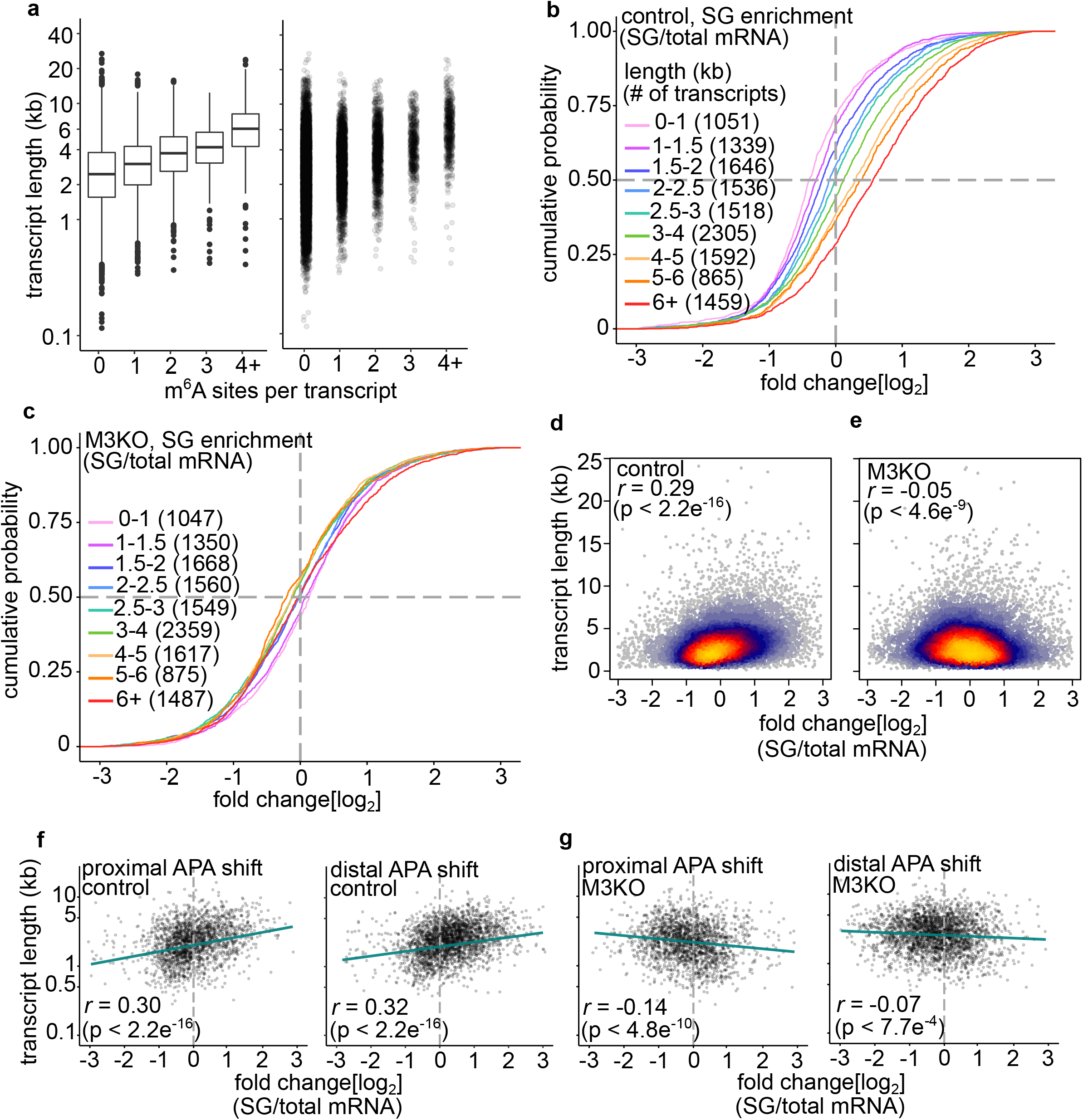
Effects of m^6^A loss and mRNA length on stress granule mRNA abundance. **a,** Transcript length and the number of m^6^A sites in a transcript are positively correlated. The left panel shows boxplots depicting the median, lower and upper quartiles of transcript length for mRNAs with an increasing number of mapped m^6^A sites. The right panel shows a scatter plot of the same data. **b,** Cumulative distribution of log_2_ fold change for mRNAs in stress granules compared to total mRNAs in control cells. A strong length-dependent trend of mRNA enrichment in control stress granules is observed. **c,** Cumulative distribution of log_2_ fold change for mRNAs in stress granules compared to total mRNAs in *Mettl3* KO cells. The length-mediated effect of stress granule mRNA enrichment is absent in *Mettl3* KO. **d,e,** Correlation between mRNA length and log_2_ fold change enrichment in stress granules relative to total mRNA levels. Panel **d** shows the result for control cells, indicating a positive correlation (r = 0.28) between length and log2 fold change enrichment in stress granules. Panel **e** shows the result for *Mettl3* KO cells, showing little correlation (r = -0.05) between length and log2 fold change enrichment of mRNA in stress granules. **f,** Proximal or distal polyadenylation site preferences do not substantially alter the length-dependent effect of mRNA enrichment in control stress granules. The length of the longest transcript isoform is plotted (y-axis) against the log_2_ fold change in stress granules relative to total mRNA (x-axis). The effect is quantified for both ‘proximal APA’ transcripts (left panel, 1960 transcripts) and ‘distal APA’ transcripts (right panel, 2612 transcripts) as determined by the analysis in Extended Data Fig. 4e. The fit line in green demonstrates a highly similar length-dependent enrichment for transcripts, irrespective of changes in APA site selection. This indicates that proximal or distal shifts in polyadenylation site usage potentially attributable to m^6^A do not substantially alter transcript enrichment in stress granules. **g,** Proximal or distal polyadenylation site preferences do not substantially alter the length-dependent effect of mRNA enrichment in *Mettl3* KO stress granules. Analysis is performed as shown in **f**, except lengths are now plotted against the enrichment of mRNAs in *Mettl3* KO stress granules. In contrast to **f**, no clear relationship can be observed between stress granule enrichment for transcripts that show either distal or proximal APA site shifts in control cells.

We therefore measured the role of mRNA length on stress granule enrichment in cells that lack m^6^A. To test this, we grouped mRNAs into bins based on their transcript length and quantified mRNAs in stress granules relative to total cellular mRNA. As expected, we observed that stress granule enrichment was markedly linked to transcript length in control MEFs (**Fig. 4b, Extended Data Fig. 4a**). The longer the transcript, the more likely it was to be enriched in stress granules. In general, transcripts greater than 4 kb were enriched in stress granules, while transcripts less than 4 kb showed de-enrichment, consistent with previous studies^6^.

Strikingly, this length-dependent mRNA enrichment is absent in the mRNAs from stress granules obtained from *Mettl3* knockout cells (**Fig. 4c, Extended Data Fig. 4b**). This suggests that many long mRNAs need m^6^A to be preferentially enriched in stress granules.

To visualize the length effect in a different way, we directly plotted mRNA length as a function of stress granule enrichment. In stress granules from control MEFs, we observed a positive correlation between mRNA length and stress granule enrichment (**Fig. 4d**). We also observed this correlation when we examined stress granule-enriched mRNAs based on previously published stress granule transcriptomes (**Extended Data Fig. 4c,d**)^6,7^. However, when we examined mRNAs enriched in stress granules from *Mettl3* knockout cells, we found no correlation between length and stress granule enrichment (**Fig. 4e**). This suggests that the correlation between transcript length and stress granule enrichment is mediated in large part by m^6^A.

### mRNA shortening does not account for the loss of the length effect in *Mettl3* knockout cells

We next considered the possibility that the reduced enrichment of normally methylated transcripts in stress granules from *Mettl3* knockout cells was due to shortening of these transcripts in these cells. It was previously found that m^6^A-containing transcripts tend to choose distal polyadenylation sites^25,39^. In a previous analysis of a subset of 661 m^6^A-containing transcripts, 70 transcripts showed use of proximal polyadenylation sites after knockdown of METTL3^39^. Although this is a small effect, one interpretation of our observations could be that METTL3 depletion causes selection of more proximal polyadenylation sites, thus leading to mRNA shortening. This could potentially reduce their enrichment in stress granules in *Mettl3* knockout cells due to decreased length, rather than decreased m^6^A.

To test this, we analyzed alternative polyadenylation site (APA) selection in both control and *Mettl3* knockout cells to determine if m^6^A altered transcript lengths. To predict APA site usage, we analyzed RNA-seq data from control and *Mettl3* knockout cells using DaPars, a program designed to measure dynamic changes in APA site usage between samples^40^. Consistent with earlier studies^39^, we found a slight overall preference for distal polyadenylation sites on transcripts in control MEFs compared to *Mettl3* knockout cells (**Extended Data Fig. 4e**). We also examined the length difference between proximal and distal polyadenylation sites to determine the effect this might have on overall transcript length. The genomic distance between alternative sites is less than 500 base pairs for most (∼70%) transcripts, and the median distance change in APA sites for all transcripts is ∼272 base pairs (**Extended Data Fig. 4f,g**). Overall, these data indicate that *Mettl3*-dependent shortening or lengthening of transcripts is relatively small but could partially account for the reduced length-dependent mRNA enrichment in stress granules in *Mettl3* knockout cells.

To determine if shifts in polyadenylation sites may have affected transcript recruitment to stress granules, we separately examined transcripts that were predicted to shift to either proximal or distal polyadenylation sites in control cells relative to *Mettl3* knockout cells (see **Extended Data Fig. 4e**). mRNAs with m^6^A-dependent shifts to either proximal or distal APA sites show similar length-dependent enrichment in stress granules in control cells (**Fig. 4f**). However, when we examined *Mettl3* knockout cells, both populations showed a loss of length-dependent enrichment in stress granules (**Fig. 4g**). Overall, this suggests that transcript length changes due to altered APA site usage cannot explain the lack of length-dependent mRNA enrichment observed in stress granules from *Mettl3* knockout cells.

Since m^6^A is often associated with the use of a distal polyadenylation site, we performed our analysis thus far using the longest isoform for each gene when determining the effects of m^6^A. However, we wanted test if this assumption by seeing if the effect would change if we examined only the shortest isoforms for each gene. Indeed, we observed that all the significant overall trends and effects were similar when we annotated mRNAs lengths based on the shortest 3’UTR isoform (**Extended Data Fig. 5a-d**). Thus, our finding that m^6^A mediates length-dependent enrichment of mRNA into stress granules is unlikely to be caused by using incorrect transcript length annotations based on the longest 3’UTR isoform, since these effects are seen regardless of the isoform annotation that is used.

It was suggested in the original study implicating mRNA transcript length in stress granule enrichment that the length of the CDS contributed disproportionately to the enrichment of transcripts in stress granules relative to the 5’UTR and 3’UTR^6^. Upon further analysis of this data, we noticed discrepancies between the sum of the transcript region lengths (5’ UTR + CDS + 3’UTR) and the total transcript lengths used in this study. We re-annotated the transcript region lengths in this dataset and found that CDS lengths were largely unchanged (**Extended Data Fig. 6 a,b**), but 3’UTR lengths for many transcripts were reduced in the original study (**Extended Data Fig. 6 c**). Adjusting for this discrepancy led to a substantial increase in the correlation between 3’UTR length and mRNA enrichment in stress granules (**Extended Data Fig. 6d**). Notably, long 3’UTRs appear to induce m^6^A methylation in mRNA^39^. Thus, both CDS and 3’UTR length correlate with m^6^A levels and stress granule enrichment, supporting the idea that length correlates with stress granule enrichment due to the ability of long exons and 3’UTRs to induce m^6^A formation in mRNA.

Since our data suggest that m^6^A depletion results in a loss of long mRNAs from stress granules, we wanted to know what might replace these mRNAs. To test this, we measured the abundance of mRNAs in stress granules using RPKM. We binned RNAs based on length, and quantified the abundance of mRNAs for each pool in stress granules and in the total cellular RNA pool (**Extended Data Fig. 7a,b**). As expected, stress granules in control cells contain fewer short and more long transcripts than their overall fractional abundance in the total cellular RNA pool. The shortest mRNAs were only ∼37% of stress granule mRNAs while they are ∼44% of total cellular mRNA, consistent with their de-enrichment in stress granules. In contrast, stress granules from *Mettl3* knockout cells were markedly different, with transcript abundance in stress granules being more similar to their abundance in the total cellular mRNA pool. Notably, the small mRNAs were no longer de-enriched and represented ∼43% of all cellular RNAs in stress granules. This suggests that the reduction of longer mRNAs in *Mettl3* knockout stress granules is compensated in part by short, abundant transcripts that make up the bulk of the total transcriptome (**Extended Data Fig. 7c**).

### m^6^A promotes length-independent enrichment of mRNAs in stress granules

Although mRNA length and stress granule enrichment correlate generally, not all mRNAs follow this trend^6^. For example, some short mRNAs are enriched while some long mRNAs are not. We next asked if m^6^A may predict enrichment irrespective of mRNA length. To test this, we first analyzed the association between transcript length, m^6^A, and stress granule enrichment using previous datasets. We found that m^6^A was a strong predictor of stress granule enrichment in U2OS cells (**Fig. 5a,b**) and NIH3T3 cells (**Extended Data Fig. 8a,b**) regardless of transcript length. This effect was the most dramatic in short transcripts, which are generally de-enriched in stress granules, but were substantially enriched when containing increasing numbers of m^6^A sites. This provides strong evidence that DF-m^6^A interactions can “override” the limitations placed on short transcripts by an RNA-RNA interaction model of mRNA recruitment to stress granules as well as substantially enhance the enrichment of longer mRNAs.

**Fig. 5:**
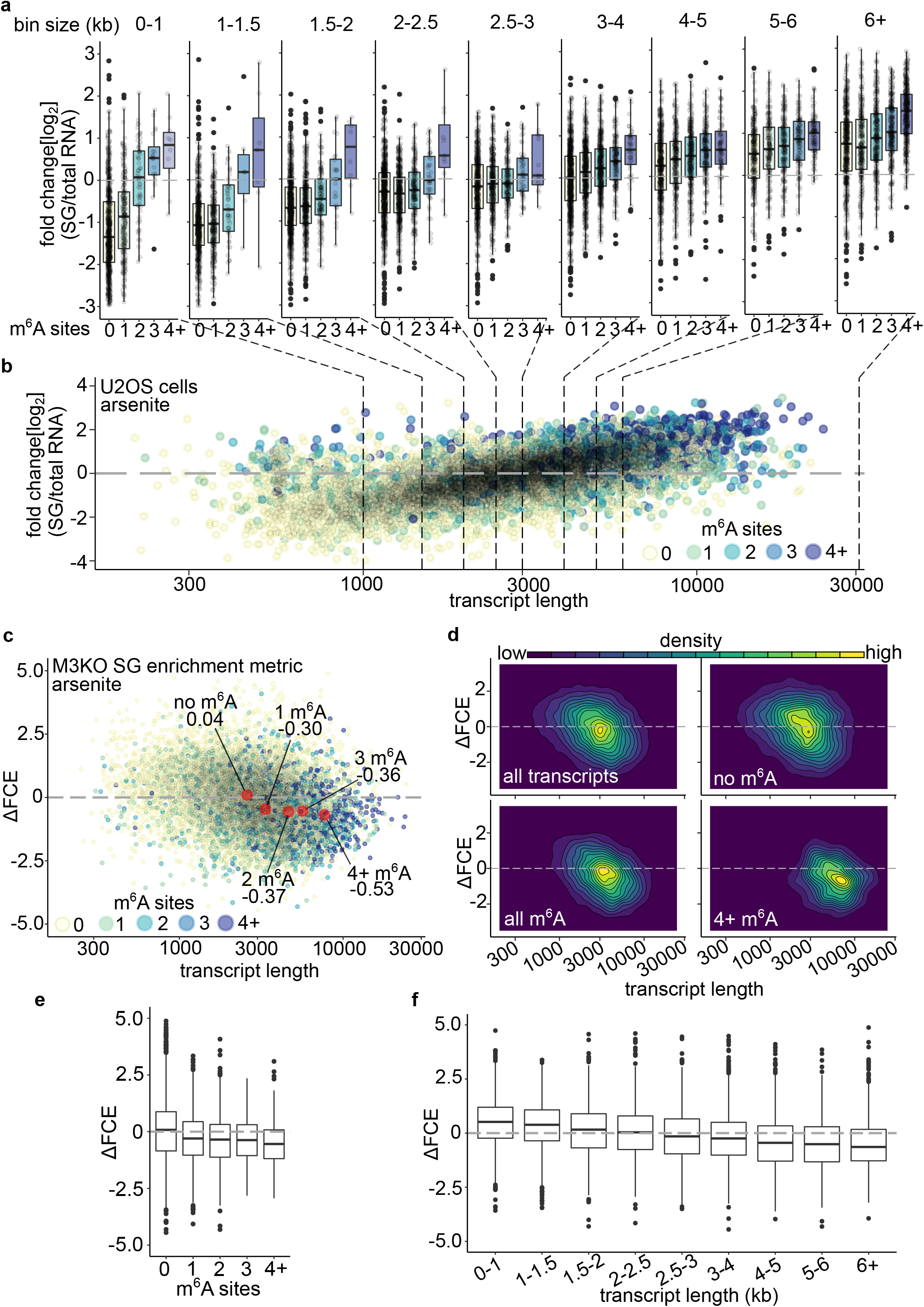
The effects of m^6^A on mRNA enrichment in stress granules is length-independent. **a,** Boxplots demonstrating the correlation between m^6^A levels, transcript length and stress granule enrichment in U2OS cells. The y-axis shows the relative level of enrichment of transcripts in stress granules relative to total cellular mRNA. The x-axis shows the number of m^6^A sites per transcript. Boxplots depict the median, upper and lower quartiles. Each dot represents a unique individual transcript. At the top of each boxplot the range of transcript lengths contained in the plot below is shown. Dotted lines at the bottom of the plot show the segment of data from the scatter blot below in **b** that is summarized. Transcript size increases from left to right. Generally, regardless of transcript length, m^6^A has a cumulative effect on the likelihood of enrichment in stress granules. **b,** Scatter plot demonstrating the correlation between m^6^A levels, transcript length and stress granule enrichment in U2OS cells. The y-axis shows the relative level of enrichment of transcripts in stress granules relative to total cellular mRNA. The x-axis shows the length of individual transcripts on a logarithmic scale. The number of m^6^A sites in each individual transcript is color-coded as shown (yellow = 0, light green = 1, blue-green = 2, blue = 3, violet = 4+). Dotted lines are shown connected to the boxplots in panel **a** that summarize the scatter data per m^6^A site. **c,** The m^6^A-dependent difference in fold-change enrichment (ΔFCE) links m^6^A, transcript length, and stress granule enrichment. ΔFCE was determined by calculating the absolute difference in log_2_ fold change enrichment between *Mettl3* KO and control stress granules relative to total mRNA for each transcript. Thus, positive ΔFCE indicates an increase in stress granule enrichment upon loss of m^6^A, while negative ΔFCE indicates a decrease in stress granule enrichment upon loss of m^6^A. Each dot represents an individual transcript’s length (x-axis) and ΔFCE (y-axis). Since differences in transcript lengths in control and *Mettl3* KO are assumed to be minor in most cases (see Extended Data Fig. 4e-g), ΔFCE reflects changes in stress granule enrichment that are largely m^6^A-dependent but length-independent. The number of m^6^A sites in each individual transcript is color-coded as shown (yellow = 0, light green = 1, blue-green = 2, blue = 3, violet = 4+). Each enlarged red dot represents the mean length and ΔFCE for all transcripts containing either 0, 1, 2, 3, or 4 or more m^6^A sites. Non-methylated mRNAs are not substantially affected, as m^6^A contributes to only ∼4% of the average transcript’s enrichment in stress granule. Low-to-moderate methylation (1-3 sites) contributes to ∼30-35% of stress granule enrichment in the average transcript, while high methylation (4+ sites) contributes to a more substantial mean of ∼50% to stress granule enrichment in the average transcript. **d,** Density plots of ΔFCE in transcripts as a function of length. The data summarizes subsets of transcripts depicted in **c**. In the top left panel, a density plot of ΔFCE from all transcripts depicted is shown. In the top right panel, a density plot of ΔFCE for only transcripts with no m^6^A is shown. Since the central mass lies at “0”, this demonstrates a negligible m^6^A-linked effect on enrichment of these transcripts in *Mettl3* KO cells. In the bottom left panel, a density plot of ΔFCE for all m^6^A-containing transcripts is shown. Since the central mass skews negative, this shows that m^6^A-containing transcripts are generally depleted in *Mettl3* KO stress granules. The bottom right panel shows a density plot of ΔFCE for only highly methylated (4+ m^6^A sites) transcripts. The majority of the density is well below “0”, indicating a strong m^6^A-linked effect on transcript depletion in *Mettl3* KO stress granules. **e,** Boxplots of ΔFCE as a function of m^6^A abundance. Boxplots depict the median and upper and lower quartiles. Loss of m^6^A substantially affects ΔFCE in the majority of m^6^A-containing transcripts. **f,** Boxplots of ΔFCE as a function of transcript length. Boxplots depict the median and upper and lower quartiles. Loss of m^6^A results in substantial increases in ΔFCE of shorter transcripts and substantial decreases of ΔFCE in longer transcripts. This shows that the enrichment of many long mRNAs is dependent on m^6^A.

We next wanted to devise a method to estimate the effect size of m^6^A on stress granule enrichment based on the length of transcripts using the *Mettl3* knockout cell stress granule transcriptome. We developed a metric called ‘Delta fold change enrichment’ (ΔFCE) which determines the overall difference in log_2_-fold change in stress granule enrichment for the same transcript in control vs. *Mettl3* knockout cells. By matching each transcript and changing only the variable of whether cells contain m^6^A, we can estimate the effect that m^6^A has on stress granule enrichment when transcript length remains relatively constant. A positive ΔFCE value for a transcript indicates a higher enrichment in stress granules in *Mettl3* knockout cells relative to control (low m^6^A-dependence), while a negative ΔFCE value indicates lower enrichment in stress granules in *Mettl3* knockout cells relative to control (high m^6^A-dependence).

Plotting this metric along with transcript length demonstrates a correlation between negative ΔFCE, transcript length, and the number of m^6^A sites (**Fig. 5c**). Based on the average effect size of m^6^A on stress granule enrichment in *Mettl3* knockout cells, ΔFCE predicts that m^6^A can increase the stress granule recruitment efficiency of a transcript by anywhere from ∼30-50% on average. Density plots which divide transcripts into groups according to their m^6^A content also demonstrate these effects (**Fig. 5d**). For transcripts with no m^6^A, the loss of m^6^A has little effect on ΔFCE, as expected. However, for transcripts with m^6^A, and especially for transcripts with numerous m^6^A sites (4+), the effects of losing m^6^A have a substantial negative effect on ΔFCE, indicating the recruitment of these mRNAs to stress granules strongly depends on m^6^A.

Grouping transcripts by m^6^A or by length also demonstrates the strong correlation between these variables and negative ΔFCE, and that greater lengths and number of m^6^A sites in a transcript corresponds to larger m^6^A-dependent effects on stress granule enrichment (**Fig. 5e,f**). Taken together, these data demonstrate that m6A can enhance enrichment of mRNAs in stress granules in a length-independent manner.

## DISCUSSION

RNA is a major component of diverse biological condensates, including stress granules. Although all cellular RNAs are found to some degree in stress granules, there appears to be factors that cause mRNA to be enriched into a stress granule relative to the overall abundance of the mRNA in the cell. Most notably, mRNA length appears to drive mRNA localization in stress granules^6^. The ability of long mRNAs to localize to stress granules has been proposed to reflect RNA self-assembly in which inter-molecular RNA-RNA interactions facilitate condensate formation^9^. Here we show that the length-dependent enrichment of mRNAs in stress granules is instead mediated in large part by m^6^A. m^6^A is preferentially targeted to long mRNAs due to the ability of long exons to induce methylation of mRNA. These m^6^A-rich regions form scaffolds for multivalent DF protein interactions that allow m^6^A mRNA-DF complexes to translocate to stress granules. Notably, the enrichment of mRNAs based on length is substantially reduced in *Mettl3* knockout cells without m^6^A. Our analysis reveals that m^6^A shapes the stress granule transcriptome by promoting the enrichment of long mRNAs in stress granules.

Stress granules are often considered a prototype for the study of mRNA-protein molecular condensates that assemble via liquid-liquid phase-separation mechanisms^2^. Thus, the mechanisms that determine which mRNAs are recruited to stress granules are of high interest due to their purported ability to reveal general principles of granule assembly. The idea that RNA-RNA interactions contribute to this process has created interest in understanding the physical principles that regulate these interactions. However, our data indicate that if RNA-RNA interactions have a role, it is unlikely to solely determine length-dependent mRNA enrichment. Although RNA-RNA interactions are important for in vitro RNA granule assembly, the conditions in vitro do not appear to fully predict RNA enrichment in cells. It should be noted that some RNA-RNA interactions clearly occur in cells, e.g. with trinucleotide and hexanucleotide repeat sequences^41^. Thus, it is possible that RNA-RNA interactions could contribute in other ways to stress granule properties.

The challenge in studying the role of mRNA length is that mRNA length is related to m^6^A levels^15,21,22^. The presence of m^6^A is not simply due to a model in which a longer mRNA simply has more opportunity to be methylated. Instead, long exons promote recruitment of the m^6^A writer complex as a result of unique histone modifications that accumulate over these exons and recruit the methyltransferase needed for m^6^A formation^23–25^. Since long exons will also lead to overall longer mRNAs, m^6^A is typically associated with long mRNAs. For this reason, some mRNAs can be long and have little m^6^A if the gene lacks these exon features. Indeed, we found that many long mRNAs that were paradoxically not highly enriched in stress granules, were simply not methylated. Thus, the discrepancy between the long RNAs that are not enriched can now be explained by the fact that it is not length per se, but m^6^A that is the major driver of length-dependent enrichment. Similarly, we can now explain the presence of the few short mRNAs that are paradoxically enriched in stress granules—these mRNAs are highly methylated.

Our data presented here, in which we depleted m^6^A and assessed changes in the stress granule transcriptome, provides a straightforward approach to remove the involvement of m^6^A in stress granule partitioning of mRNA, without impairing putative RNA-RNA interactions or other aspects of RNA self-assembly. Since the length-dependent enrichment of mRNAs in stress granules is markedly reduced in *Mettl3* knockout cells, the role of RNA self-assembly is likely to be minor relative to m^6^A, except in special cases. Notably, the overall abundance of very long transcripts (5+ kb) in *Mettl3* knockout stress granules was slightly greater than the overall abundance in total cellular mRNAs (see **Extended Data Fig. 7b**). This suggests that RNA-RNA interactions may play a role in RNA recruitment or retention of especially long transcripts in stress granules, even if the primary mechanism that links transcript length to stress granules for the majority of transcripts is m^6^A.

A recent study looked at methylated transcripts in control cells and “ΔMETTL3” mouse embryonic stem cells to examine the effect of m^6^A on mRNA enrichment in stress granules^42^. In this study, fluorescence in situ hybridization (FISH) showed that three m^6^A-containing mRNAs showed little difference in their stress granule enrichment levels in wild type or ΔMETTL3 cells. This study was problematic because the ΔMETTL3 cell line contains a truncated METTL3 isoform and is not in fact depleted of m^6^A, limiting the usefulness of these cells when studying the full effects of m^6^A on mRNA^26^. Nevertheless, our study highlights that the critical function of m^6^A is not to determine whether a transcript is localized in a stress granule or not, but whether m^6^A confers additional enrichment beyond the overall concentration of the mRNA in the cytoplasm. The use of full *Mettl3* knockout cells, combined with an analysis of all methylated transcripts provides a much more sensitive indicator of the role of m^6^A rather than looking at a small number of transcripts in individual cells. Additionally, a transcriptomic analysis is more useful than FISH to compare concentration of mRNA in a stress granule compared to the cytosol compared, in which accessbility of the probe into the granule is likely different than in the cytosol. Overall, the results here provide a comprehensive assessment of the role of m^6^A in targeting mRNAs to stress granules.

The number of m^6^A sites in an mRNA is linked to the degree of stress granule enrichment of that mRNA. The number of m^6^A sites is important since it will determine the number of bound DF proteins, which will in turn lead to greater partitioning of DF-mRNA complexes in stress granules.^11^. m^6^A thus facilitates proximity-induced multivalent interactions between DF proteins, and these interactions are most likely to occur on transcripts containing long exons that are rich in m^6^A. Our data shows these interactions are critical for shaping the stress granule transcriptome and controlling length-dependent enrichment of mRNAs.

## METHODS

### *Mettl3* knockout mouse embryonic fibroblasts

Mouse embryonic fibroblasts were generated from *Mettl3^flox/flox^* mice, a generous gift from the lab of Michael Kharas. The generation of these mice has been described previously^31^. In brief, *loxP* sites were inserted that spanned the fourth exon of *Mettl3*. Homozygous *Mettl3^flox/flox^*embryos were harvested at E13.5. The embryos were separated mechanically and treated with trypsin. The separated cells were then plated and grown for three passages. After the third passage, the cells were transduced with SV40 large T-antigen lentivirus and passaged until they reached senescence. Cells that escaped senescence were than transduced with the Cre-ERT2 lentivirus and selected by puromycin treatment. Single-cell clones were then isolated from the surviving cells. Cells were then treated with 500 nM 4-hydroxytamoxifen (4-OHT) for 48 h to induce activation of Cre recombinase. After 7 days, the complete loss of METTL3 was confirmed by western blot, and the complete loss of m^6^A was confirmed using thin-layer chromatography.

### Cell culture

Mouse embryonic fibroblasts (MEFs) were grown in Dulbecco’s Modified Eagle Medium (DMEM) containing 10% fetal bovine serum (FBS) with 100 U ml-1 of penicillin and 100 ug mL-1 of streptomycin under standard culturing conditions (37°C with 5.0% CO2). Cells were passaged as required using trypsin according to manufacturer’s instructions.

### Cellular stress

Cellular stresses were performed as follows: heat shock at 42°C for by partial submersion in a water bath for 30 min; arsenite stress by addition of DMEM with 0.5 mM of sodium arsenite for 30 min; thapsigargin stress by addition of DMEM with 1 uM of thapsigargin for 1 h; sorbitol stress by addition of DMEM with 400 mM sorbitol for 2 h.

### 2-dimensional thin layer chromatography

2-dimensional thin layer chromatography was performed as previously described^43^. In brief, twice-purified polyadenylated mRNA was digested with T1 ribonuclease. This results in mRNA cleavage immediately following guanosine residues. The RNA mixture is then labeled with [γ-^32^P]ATP using T4 PNK, so that all nucleotides following a guanosine are labelled with ^32^P on the 5’ end. Excess [γ-^32^P]ATP is removed by apyrase. RNA is then digested into single nucleotides using P1 nuclease. The digested RNA was then spotted on PEI-cellullose plates and developed as previously described^43^. Radioactively labelled nucleotides were detected using a phosphor storage screen and Amersham Biosciences Typhoon 9400 Variable Mode Imager.

### Immunostaining

Cells were plated to reach confluency of 40-60% on the following day in glass-bottom well plates. Cells were gently washed with 1xPBS twice to remove media and serum proteins. Cells were then fixed with 4% paraformaldehyde in 1xPBS for 10 min. Fixed cells were washed three times with 1xPBS and incubated with blocking solution (3% FBS, 0.2% Triton X-100 in 1xPBS) for 30 min. Cells were then incubated with primary antibody diluted 1:100 in blocking solution for 60 min. Cells were then washed three times with 1xPBS and incubated with secondary antibody at 2 µg ml^-1^ in blocking solution for 60 min. Cells were then washed once with 1xPBS and then incubated with Hoechst stain at 0.1 µg ml^-1^ in ddH_2_O for 10 min. Cells were then washed one final time with ddH_2_O. The cells were then mounted with coverslips using Prolong Diamond Antifade Mountant. All steps were carried out at 25°C.

### Western blotting

Cells were lysed following growth to confluency using RIPA buffer (10 mM Tris-HCl pH 7.4, 10 mM EDTA, 50 mM NaF, 50 mM NaCl, 1% Triton X-100, 0.1% SDS) and shaken vigorously at 4°C for 45 min. Insoluble material was spun down at 4°C and the supernatant was collected. Protein was quantified using the BCA assay. Equal quantities of protein were loaded in 4-12% Bis-Tris gels and were separated by constant 140 V for 90 min. Separated proteins were then transferred to nitrocellulose membranes by constant 10 V in 4°C overnight. Membranes were washed in ddH_2_O and blocked with 3% FBS in PBS-Tween (0.2%) for 1 h. Membranes were blotted with primary antibodies diluted in blocking solution at 4°C overnight. Membranes were then washed extensively with PBS-Tween and further incubated with secondary antibodies conjugated to horseradish peroxidase (HRP) in blocking solution at 25°C for 1 h. Membranes were extensively washed in PBS-Tween before a brief incubation with ECL reagent (Amersham). Blots were imaged on the ChemiDoc XRS+ system (Bio-Rad).

### Antibodies

The following antibodies were used for immunofluorescence experiments: mouse anti-TIAR (Clone 6; 610352, BD Biosciences), rabbit anti-YTHDF2 (24744-1-AP, Proteintech), mouse anti-IgG Alexa Fluor 488 (A11001, Invitrogen), rabbit anti-IgG Alexa Fluor 594 (A11012, Invitrogen). The following antibody was used for western blotting: rabbit anti-METTL3 (15073-1-AP, Proteintech).

### Stress granule measurements

Stress granule measurements were obtained using Fiji (ImageJ). First, image color channels were separated. A black-and-white mask was created of the Hoechst-stained channel using ‘Image -> Adjust -> Threshold ‘and adjusting the settings to best image fit. These regions were then subtracted from the stress granule channel stained with TIAR to remove nuclear fluorescence. ‘Image -> Adjust -> Threshold ’was then used on the TIAR channel to obtain a mask of stress granules. The mask was refined by smoothing edges (Process -> Smooth) and performing a watershed on closely adjacent or overlapping stress granules (Process -> Binary - > Watershed). This mask was then used to create regions of interest in the ImageJ ROI manager (Analyze -> Tools -> ROI manager). Area measurements for stress granules were obtained (Analyze -> Analyze Particles). This mask was used for determining regions of interest used in downstream quantification.

### Fluorescent trace quantification

A 2.57 um line was drawn through the center of stress granules and the underlying pixel intensity values for the staining of both TIAR and YTHDF2 were recorded. This line size was chosen as it was generally large enough to span the entire diameter of medium-to-large sized stress granules. For each of these measurements, a corresponding background measurement of fluorescent intensity in the cytoplasmic region immediately outside the stress granule was taken. The average fluorescent intensity from ∼150 nm increments of the 2.57 um line from stress granules and the adjacent cytoplasm were then grouped and plotted. Representative images were used as input.

### Protein partition coefficients

Protein partition coefficients were calculated essentially as described previously^11^. In brief, the stress granule regions of interest were measured for average fluorescence intensity using ImageJ. The stress granule region of interest was then moved to the cytoplasm adjacent to the stress granule of interest and a corresponding cytoplasmic average pixel intensity was obtained. The average intensity value from the stress granule was then divided by the average intensity value from the cytoplasm. The resulting partition coefficients were then plotted. Representative images were used as input.

### Stress granule purification

Stress granule purification was performed as described previously^7^. In brief, cells grown to confluence on 15 cm dishes were subjected to stresses, spun down and snap frozen in liquid nitrogen before proceeding with purification.

### RNA-seq and differential expression analysis

For RNA-seq of stress granules, RNA was first phenol-chloroform extracted from purified stress granules^7^. Polyadenylated mRNA was then purified from total RNA using Dynabeads conjugated to oligo(dT). PolyA mRNA was then eluted from beads and quantified using a Qubit (ThermoFisher). 20 ng of polyA-purified mRNA samples were processed and sequenced by Genewiz, LLC using an Illumina HiSeq 3000.

FastQ files were checked for read quality by analysis with FastQC^44^. Adapters were removed by FLEXBAR and trimmed for read quality (phred >= 30)^45^. Before alignment to the transcriptome, all reads mapping to rRNA and tRNA were removed. After rRNA read removal, the resulting reads were aligned to the Ensembl mouse transcriptome (GRCm38.75/mm10)^46^. All alignment steps were carried out using STAR genome aligner^47^. The resulting read count tables were used as input for all downstream RNA-seq analysis.

In brief, reads per kilobase per million reads (RPKM) was calculated by determining the total number of reads mapped to a gene, multiplying by a per-gene length normalizing factor, multiplied by the sequencing depth of the library. This was then divided by the total number of mapped reads in each sample. The average of two biological replicates was used to calculate any RPKM data presented.

Differential expression analysis was performed using the R package DESeq2^48^. The output of this analysis was used to create all cumulative distribution plots and boxplots of log2fold change. The R package ggplot2 was used to create all data visualizations associated with RNA-seq data^49^.

Transcript lengths were obtained by combining the lengths of all transcribed exons of each transcript (GRCm38.75/mm10). The longest transcript isoforms were used unless explicitly stated otherwise.

The m^6^A peak-containing bed file was processed to produce a table of m^6^A peak counts per gene. RNA-seq data from the U2OS stress granule transcriptome was obtained from data in Khong A, et al. 2017^6^. RNA-seq data from the NIH3T3 stress granule transcriptome was obtained from data in Namkoong S, et al. 2018^7^. m^6^A MeRIP-seq data from E13.5 MEFs (GSE61995) were downloaded from GEO^27^. m^6^A MeRIP-seq data from U2OS cells (GSE92867) were downloaded from GEO^50^.

### Polyadenylation site analysis

Alternate polyadenylation (APA) site analysis was performed using DaPars^40^. Briefly, RNA-seq gene counts were used as input to predict changes in polyadenylation site usage in control cells relative to *Mettl3* knockout cells. Transcripts with a greater than 5% deviation in APA site usage in control relative to *Mettl3* knockout cells were defined as proximally or distally shifted, depending on the directional change in polyadenylation site usage. When mapping m^6^A sites to transcripts identified in the APA analysis, the longest transcript isoform was used.

### Statistical analysis

All statistical analyses were performed using the two-sided student’s t-test unless stated otherwise. Significance cutoffs were defined as p = 0.05. All data visualizations were created using either GraphPad Prism (9.2.0) or R (4.1.1)

### Image acquisition and analysis

All fluorescent images were collected with a wide-field fluorescent microscope (Eclipse TE2000-E, Nikon). All images were acquired using NIS-Elements Viewer software (Nikon). Images were analyzed using Fiji (ImageJ v2.10).

### Data availability

RNA-seq data and all associated analysis used for data visualization reported in this paper have been deposited in the Gene Expression Omnibus (GEO) under accession number GSE190121. All code used to perform analysis or create figures is available from the corresponding author upon reasonable request.

## Acknowledgements

We thank all members of the Jaffrey laboratory for comments and suggestions. We also thank the lab of M. Kharas for generously providing *Mettl3^flox/flox^* mice to make the cell lines used in this study. This work was supported by the National Institutes of Health grants R35NS111631 and R01CA186702 (S.R.J.), F31CA254763 (R.J.R.), and National Research Foundation of Korea grant NRF-2020R1C1C1009253 (S.N.).

## Author contributions

R.J.R. and S.R.J. conceived the project and designed the experiments. R.J.R. performed all experiments unless stated otherwise. B.F.P. isolated MEFs and established cell lines used in the study. H.X.P. performed thin-layer chromatography of m^6^A in mRNA. S.N. purified the stress granules. R.J.R. designed and prepared the figures. R.J.R. and S.R.J. wrote the manuscript. The manuscript was read and approved by all authors.

---

**Extended Data Fig. 1:**
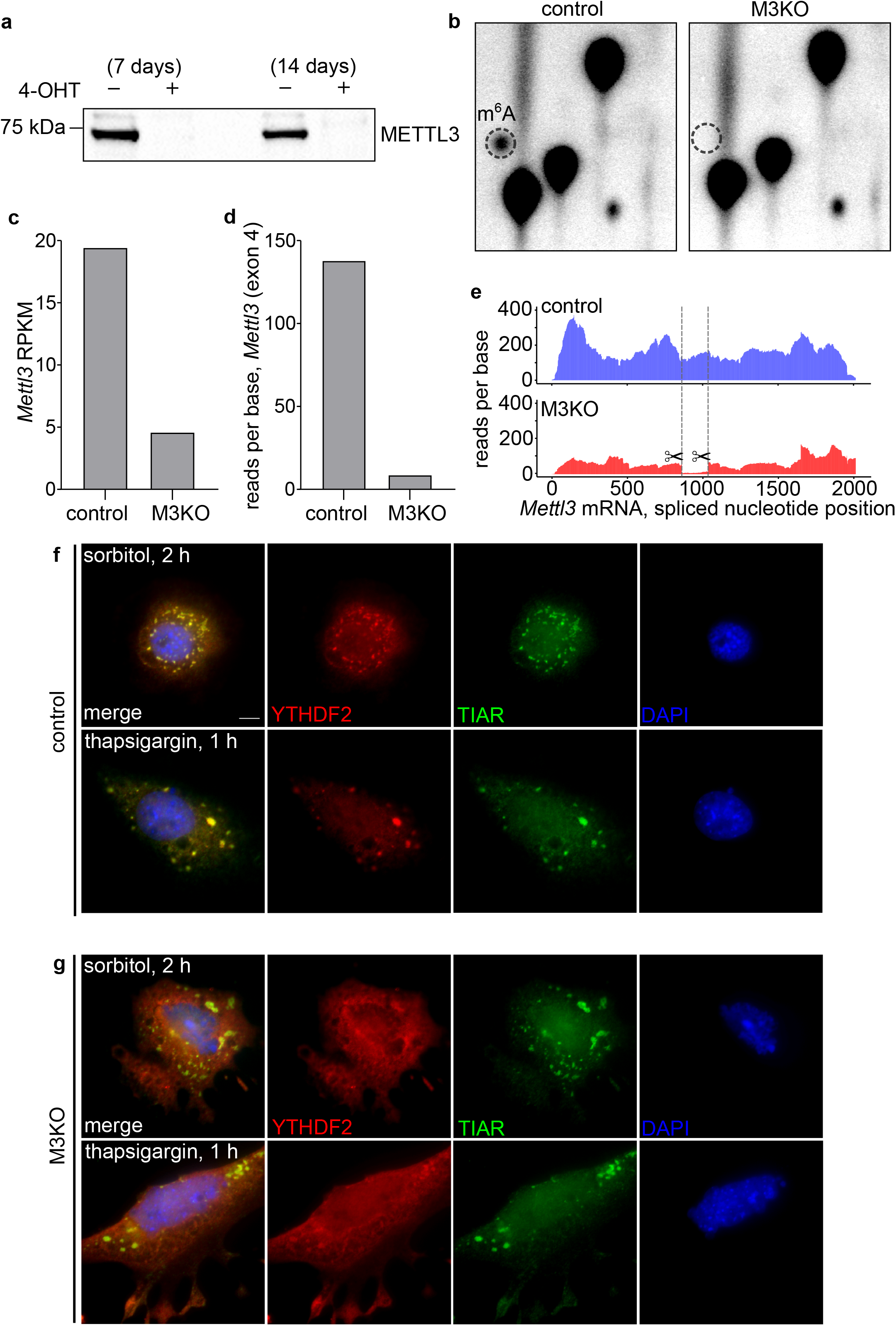
Validation of 4-OHT mediated loss of *Mettl3* and additional stress responses. **a,** *Mettl3* is depleted with high efficiency in *Mettl3 f/f* MEFs cells treated with 4-hydroxytamoxifen (4-OHT) for 48 h. The first lane shows the level of *Mettl3* in cells untreated with 4-OHT, while the second lane shows the level of *Mettl3* in the same cells seven days after treatment with 500 nM of 4-OHT. The middle lane is blank. The lasts two lanes show protein collected from cells passaged for two weeks after 4-OHT treatment. All subsequent experiments were performed between one and two weeks after 4-OHT treatment. **b,** Thin-layer chromatography of twice-polyA-purified RNA extracted from MEFs on the eighth day after treatment with 4-OHT. Control mRNA is depicted in the left panel, and mRNA from *Mettl3* knockout cells (M3KO) on the right. The dotted circle shows the position at which m^6^A migrates, demonstrating a near-complete loss of m^6^A in cells following 4-OHT treatment and loss of METTL3 protein. **c,** 4-OHT treatment results in a substantial reduction in *Mettl3* mRNA expression relative to control. Reads per kilobase per million reads (RPKM) was calculated for *Mettl3* in samples from control and *Mettl3* KO cells. Total RNA-seq reads mapping to *Mettl3* mRNA are reduced by a factor of approximately two log_2_ fold. Data reflects the averages of two biological replicates per condition. **d,** 4-OHT treatment results in a significant loss of reads mapping to exon 4 in *Mettl3* KO cells relative to control. The average number of unique reads per base was calculated for *Mettl3* exon 4 (GRCm38.p6; ENSMUSE00001262239; Chr14:52,298,640-52,298,815) which is targeted for deletion during 4-OHT-mediated Cre activation. In *Mettl3* KO cells, reads mapping to exon 4 were reduced by approximately 95%. The targeting of exon 4 is significant, because this region codes for a zinc-finger domain required for methylation activity. Data reflects the averages of two biological replicates per condition. **e,** 4-OHT treatment results in a global reduction of RNA-seq reads mapping to *Mettl3* mRNA, and a significant specific reduction of reads on exon 4. A histogram showing the reads per nucleotide (y-axis) along the length of the spliced *Mettl3* mRNA (x-axis). Control (blue) shows a substantially higher total number of mapped reads relative to *Mettl3* KO (red). The dotted lines delineate the boundaries of exon 4 within *Mettl3*. The scissors signify the Cre-mediated removal of this region after treatment with 4-OHT in *Mettl3* KO cells. This demonstrates both the substantial loss of *Mettl3* reads in *Mettl3* KO cells and the specific loss of the deleted region in exon 4. Data reflects the averages of two biological replicates per condition. **f,** Response to sorbitol and thapsigargin stress in control MEFs. Sorbitol treatment (400 mM) was performed for 2 h at 37°C. Thapsigargin treatment (1 uM) was performed for 1 h at 37°C. YTHDF2 is depicted in red, TIAR is depicted in green, and DAPI staining is depicted in blue. **g,** Response to sorbitol and thapsigargin stress in *Mettl3* KO cells. Conditions are as described in **f**.

**Extended Data Fig. 2:**
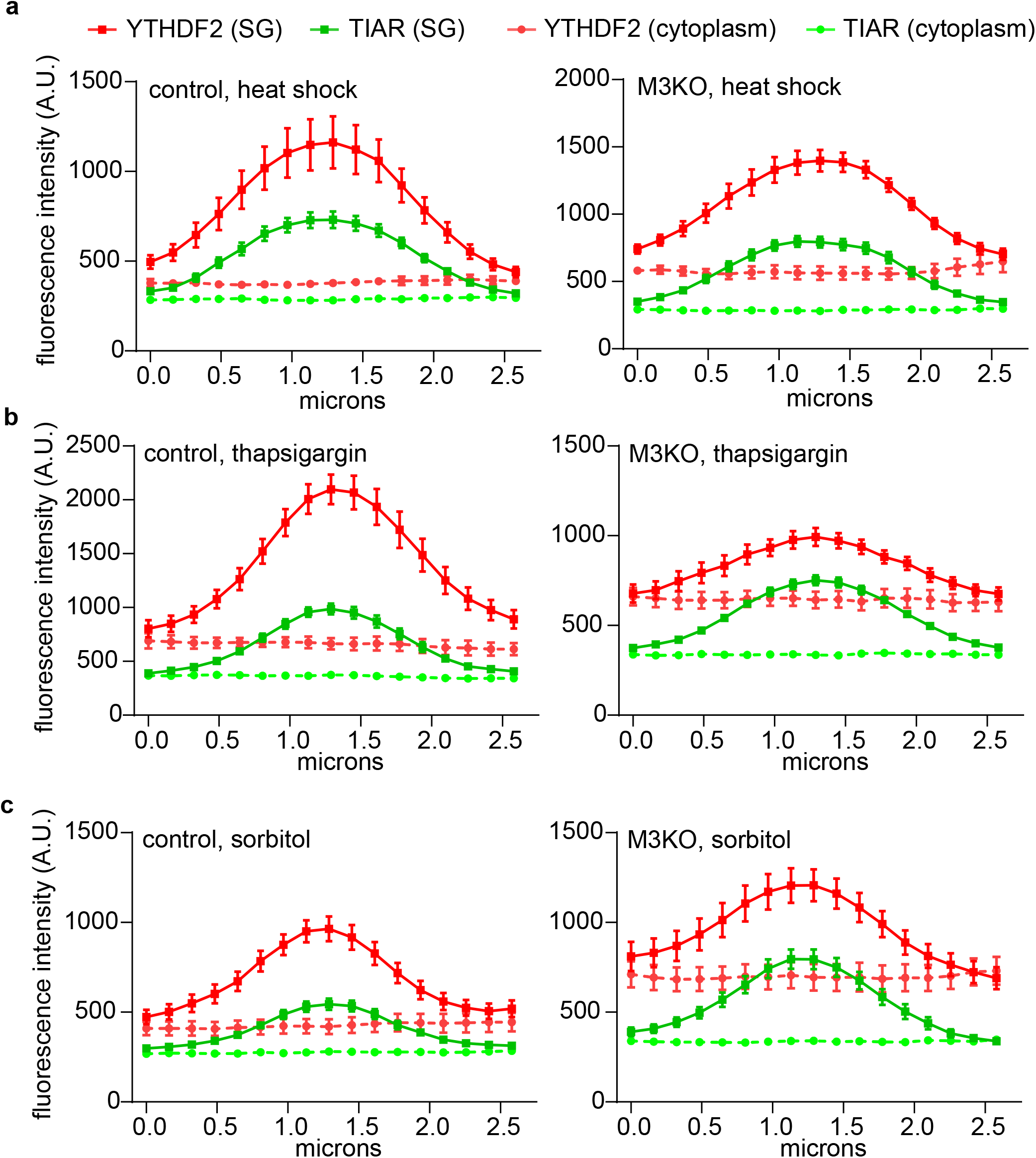
YTHDF2 and TIAR partition efficiency into stress granules. **a-c**, Quantifying changes in fluorescence intensity values in stress granules after heat shock, thapsigargin, and sorbitol treatment. The results are similar to those obtained by measuring average partition coefficients for entire granules as shown in Fig. 2c. Traces for TIAR are shown to demonstrate the corresponding bounds of the granules that were chosen for the analysis.

**Extended Data Fig. 3:**
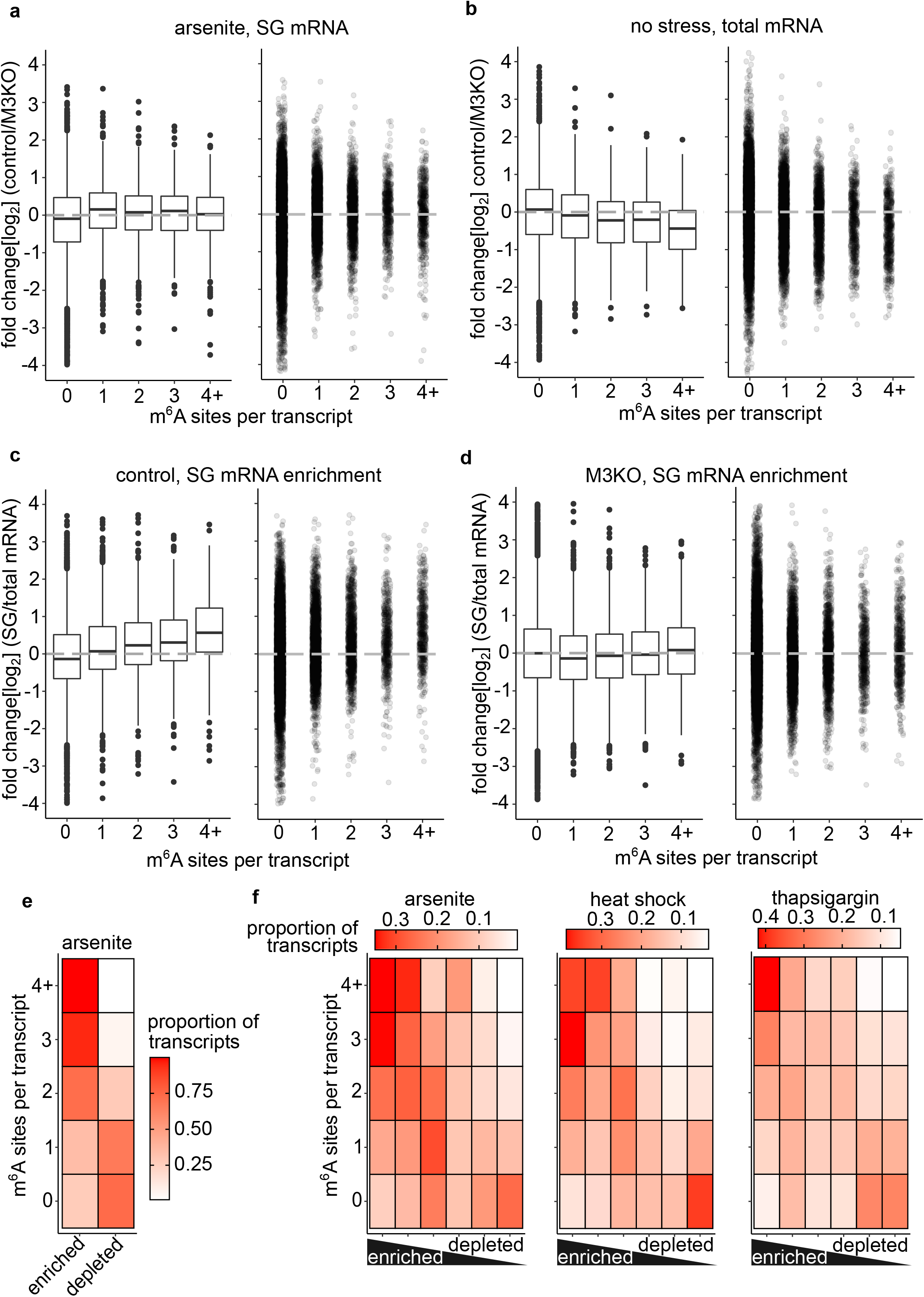
Effects of m^6^A loss on total and stress granule mRNA relative abundance. **a,** Boxplot and scatter plot of log_2_ fold change for transcripts in control and *Mettl3* KO stress-granule enriched mRNAs after arsenite treatment. Data are identical to that presented as a cumulative distribution in Fig. 3a. m^6^A-containing transcripts are slightly enriched in control compared to *Mettl3* KO. **b,** Boxplot and scatter plot of log_2_ fold change for transcripts in control and *Mettl3* KO total mRNA samples of total cellular mRNA. Data are identical to that presented as a cumulative distribution in Fig. 3b. The abundance of non-m^6^A containing mRNAs is relatively unchanged between control and *Mettl3* KO, while the abundance of m^6^A-containing transcripts is substantially reduced in control due to the destabilizing effects of m^6^A. **c,** Boxplot and scatter plot of log_2_ fold change for mRNAs in stress granules compared to total mRNAs in control cells. Data are identical to that presented as a cumulative distribution in Fig. 3c. m^6^A-containing transcripts are enriched in arsenite-induced stress granules, and the effect size increases with the number of m^6^A sites. **d,** Boxplot and scatter plot of log_2_ fold change for mRNAs in stress granules compared to total mRNAs in *Mettl3* KO cells. Data are identical to that presented as a cumulative distribution in Fig. 3d. m^6^A does not have a substantial effect on the enrichment of mRNAs in arsenite-induced stress granules. **e,** Heatmap of m^6^A transcript proportions in mRNAs enriched and depleted from arsenite-induced stress granules in U2OS cells. Data is from Khong et al. 2017. Stress granule enriched mRNAs were defined by the authors as those with a positive (+) 1.0 log_2_ fold change abundance in stress granule RNA relative to total cellular RNA. Stress granule depleted mRNAs were defined as having a negative (-) 1.0 log_2_ fold change or less. Stress granule RNA levels were normalized to total cellular RNA levels before stress. **f,** Heatmap of m^6^A transcript proportions in mRNAs enriched and depleted from arsenite, heat shock, and thapsigargin induced stress granules in NIH3T3 cells. Data is from Namkoong et al. 2018. Transcripts were organized by the authors into six clusters using a mathematical k-means clustering model with mRNAs in the first group (far left) being defined as the most enriched, and the mRNAs in the last group (far right) being defined as the most depleted in stress granules. These clusters were measured in stress granule enriched mRNAs relative to cytoplasmic mRNAs harvested at the same time point in cells relative to total mRNA before stress.

**Extended Data Fig. 4:**
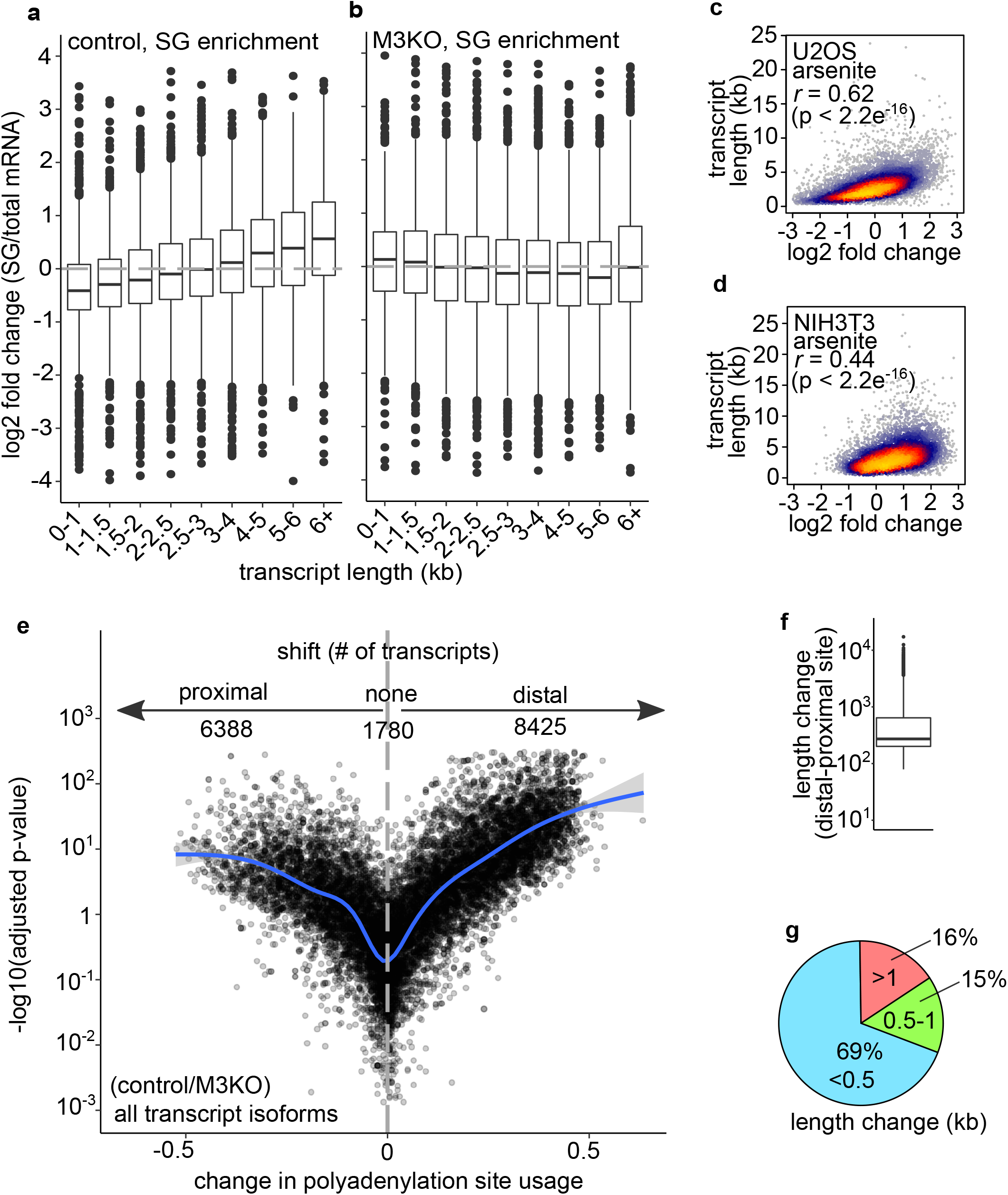
Effects of m^6^A loss on stress granule enrichment and analysis of alternative polyadenylation. **a,** Boxplot of log_2_ fold change for mRNAs in stress granules compared to total mRNAs in control cells. Data are identical to that presented as a cumulative distribution in Fig. 4b. Shorter transcripts are generally de-enriched in control stress granules, while longer transcripts are generally enriched. **b,** Boxplot of log_2_ fold change for mRNAs in stress granules compared to total mRNAs in *Mettl3* KO cells. Data are identical to that presented as a cumulative distribution in Fig. 4c. The length-dependent effect of mRNA enrichment in stress granules observed in control cells is generally absent in *Mettl3* KO. **c,** Correlation of length and mRNA stress granule enrichment in U2OS cells after arsenite treatment. Processed transcript length and log_2_ fold change data was taken from a previous analysis^6^. A strong positive correlation (r = 0.6) was observed between transcript length and stress granule enrichment of mRNAs. Notably, the protocol used to generate this dataset involves the immunoprecipitation of an endogenously-labeled G3BP1 protein during preparation of the stress granule fraction, whereas our protocol involves sequencing of the total granule fraction^7^. **d,** Correlation of length and mRNA stress granule enrichment in NIH3T3 cells after arsenite treatment. Processed log2 fold change data was taken from a previous analysis^7^. Transcript lengths were calculated from GRCm38.p6 (mm10). A positive correlation (r = 0.44) was observed between transcript length and stress granule enrichment of mRNAs. **e,** The presence of m^6^A correlates with a slight preference for distal polyadenylation site usage. We used DaPars^40^, a program for predicting changes in polyadenylation site usage from RNA-seq data, to determine if polyadenylation site usage was altered upon the loss of m^6^A. This provides an output (shown on the x-axis) that predicts proximal (negative values) or distal (positive values) shifts in polyadenylation site usage relative to another experimental condition. Each dot represents a unique transcript isoform. The -log10 p-value for each transcript is plotted along the y-axis. A best-fit line for the data is shown in blue and deviation from the fit is shown in light gray. The overall number of transcripts assigned to each category is shown at the top of the panel. Transcripts are assigned to the ‘proximal’ (6388 transcripts) and ‘distal’ (8425 transcripts) group if the change in site usage is >5% in control relative to *Mettl3* KO. Transcripts with <5% change are not assigned (1780 transcripts) to either group. A slight overall preference for distal site usage in control relative to knockout is observed both in terms of overall transcripts and likelihood of significance. **f,g,** Selection of proximal or distal polyadenylation sites has a small effect on overall transcript length. Although we did not observe a substantial effect on stress granule enrichment in Fig. 4d-e, we wanted to determine how overall length of transcripts could be affected by selection of distal or proximal polyadenylation sites. **f** shows a boxplot depicting the length change between proximal and distal polyadenylation site usage in all transcripts depicted in **e**. The median length change is approximately 272 base pairs. **e** shows that the vast majority (∼70%) of alternative polyadenylation sites will cause a length change of less than 0.5 kb in the final transcript. Only a small number of transcripts (∼16%) have an alternative polyadenylation site that will lead to a change in transcript length of greater than 1 kb.

**Extended Data Fig. 5:**
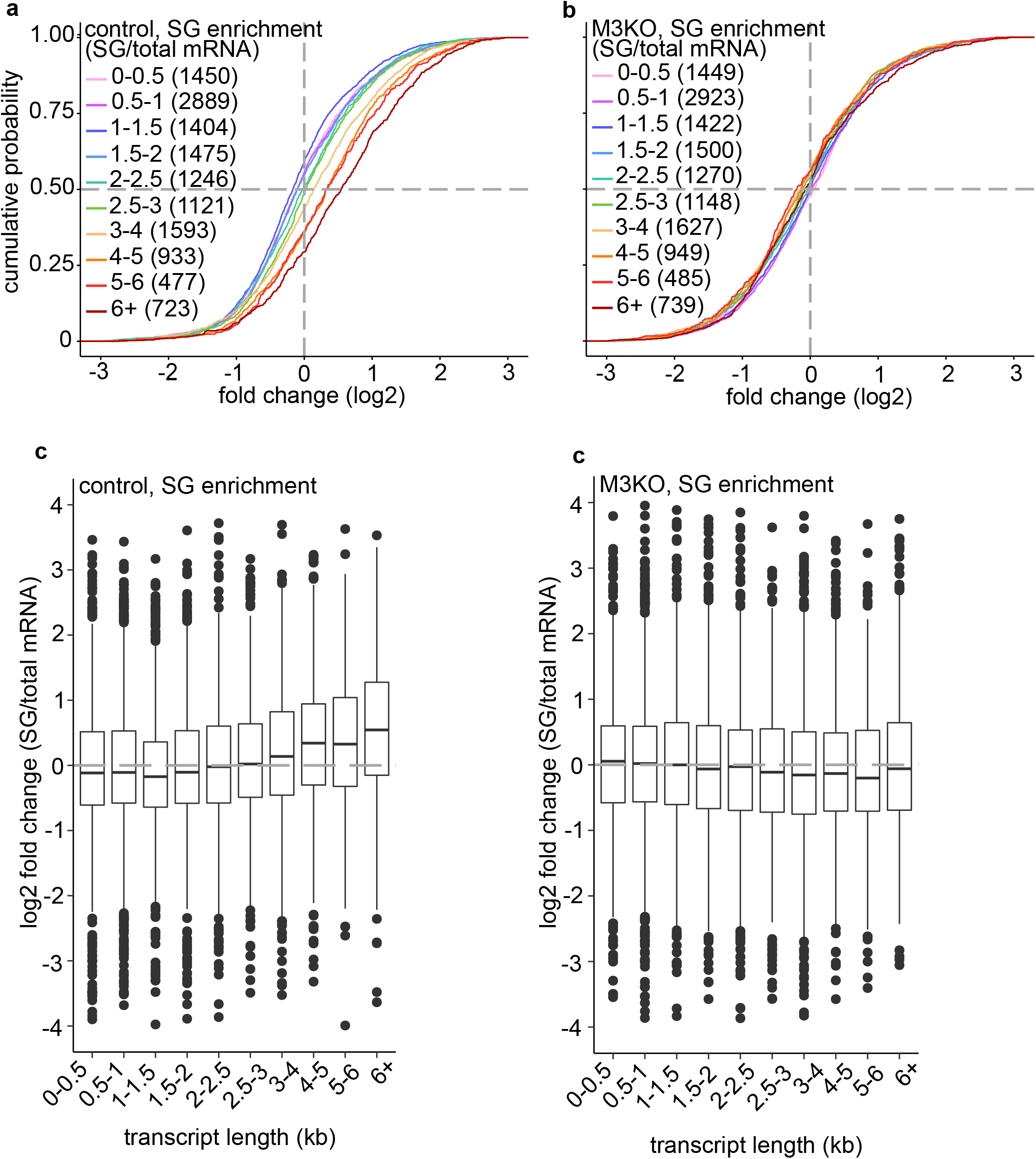
Analysis of mRNA enrichment using shortest isoforms. **a-b**, Cumulative distribution plots of differential expression using shortest isoforms. Data analysis and interpretation is essentially the same as Fig. 4b,c. **c-d**, Box plots of differential expression using shortest isoforms. Data analysis and interpretation is essentially the same as Extended Data Fig. 4a,b.

**Extended Data Fig. 6:**
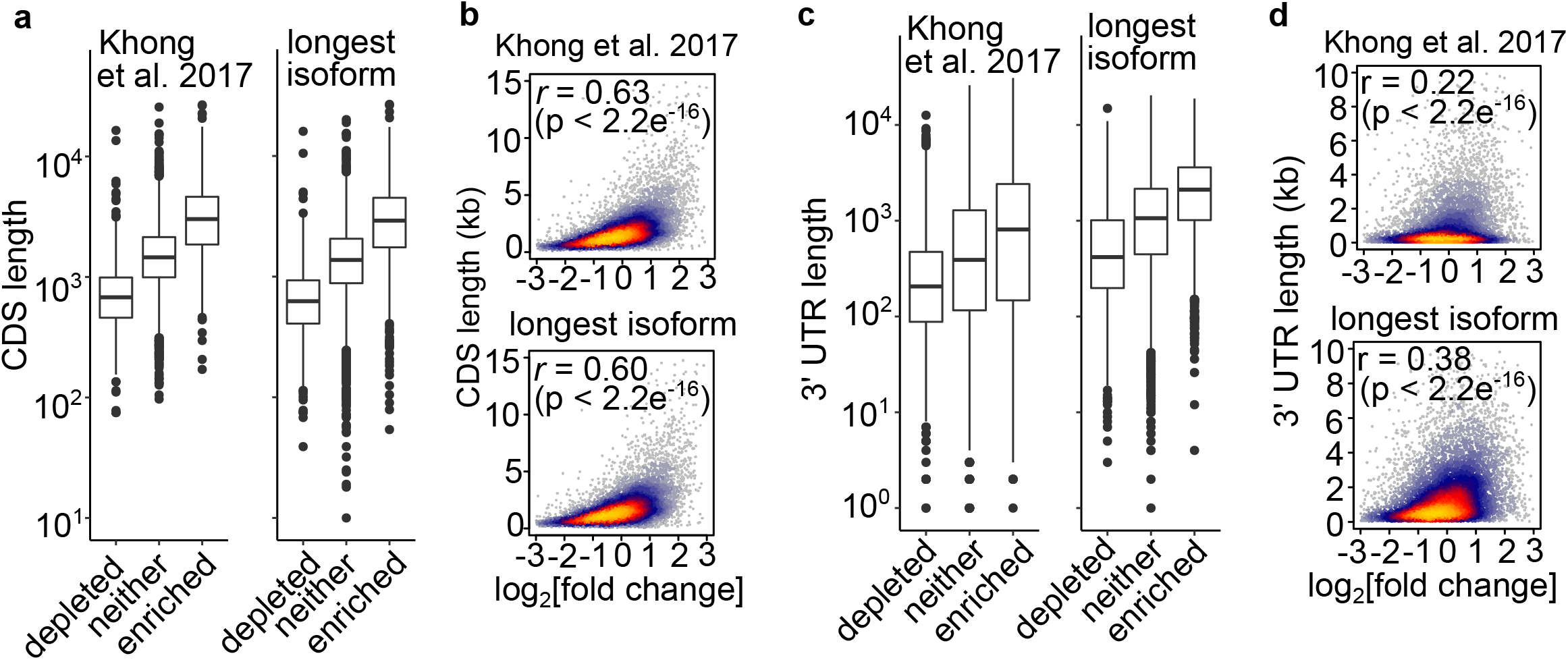
Re-analysis of the CDS and 3’UTR effect on stress granule enrichment in U2OS cells. **a**, Comparison of CDS lengths used in Khong et al. 2017 with re-annotated CDS lengths from the longest transcript isoforms in GRCh37.p13. The left panel shows the CDS lengths used in the original annotation provided by Khong et al. 2017, while the right panel shows the CDS lengths retrieved for the longest mRNA isoforms from GRCh37.p13. Stress granule enriched mRNAs were defined as having >1.0 log_2_ fold change or greater. Stress granule depleted mRNAs were defined as having <-1.0 log_2_ fold change or less. mRNAs between -1.0 and 1.0 log_2_ fold change are grouped into the ‘neither’ category. Since the dataset in Khong et al. 2017 does not contain transcript identifiers, the source of the discrepancy in lengths could not be determined. However, the re-annotation has a limited effect on CDS transcript lengths. **b**, Correlation between CDS length and enrichment of mRNAs in stress granules. Since the effects seen in **a** are relatively small, there was very little effect on the correlation between CDS and mRNA enrichment in stress granules in the original annotation (top panel, *r* = 0.63) and the re-annotation (bottom panel, *r* = 0.60). **c**, Comparison of 3’UTR lengths used in Khong et al. 2017 with re-annotated 3’UTR lengths from the longest transcript isoforms in GRCh37.p13. The left panel shows the 3’UTR lengths used in the original annotation provided by Khong et al. 2017, while the right panel shows the 3’UTR lengths retrieved for the longest mRNA isoforms from GRCh37.p13. This resulted in a substantial increase in the length of 3’UTRs in each category. **d**, Correlation between 3’UTR length and enrichment of mRNAs in stress granules. The general increase in the length of the 3’UTR had a substantial effect on the correlation between 3’UTR length and mRNA enrichment in stress granules when comparing the original annotation (top panel, *r* = 0.22) and the re-annotation (bottom panel, *r* = 0.38).

**Extended Data Fig. 7:**
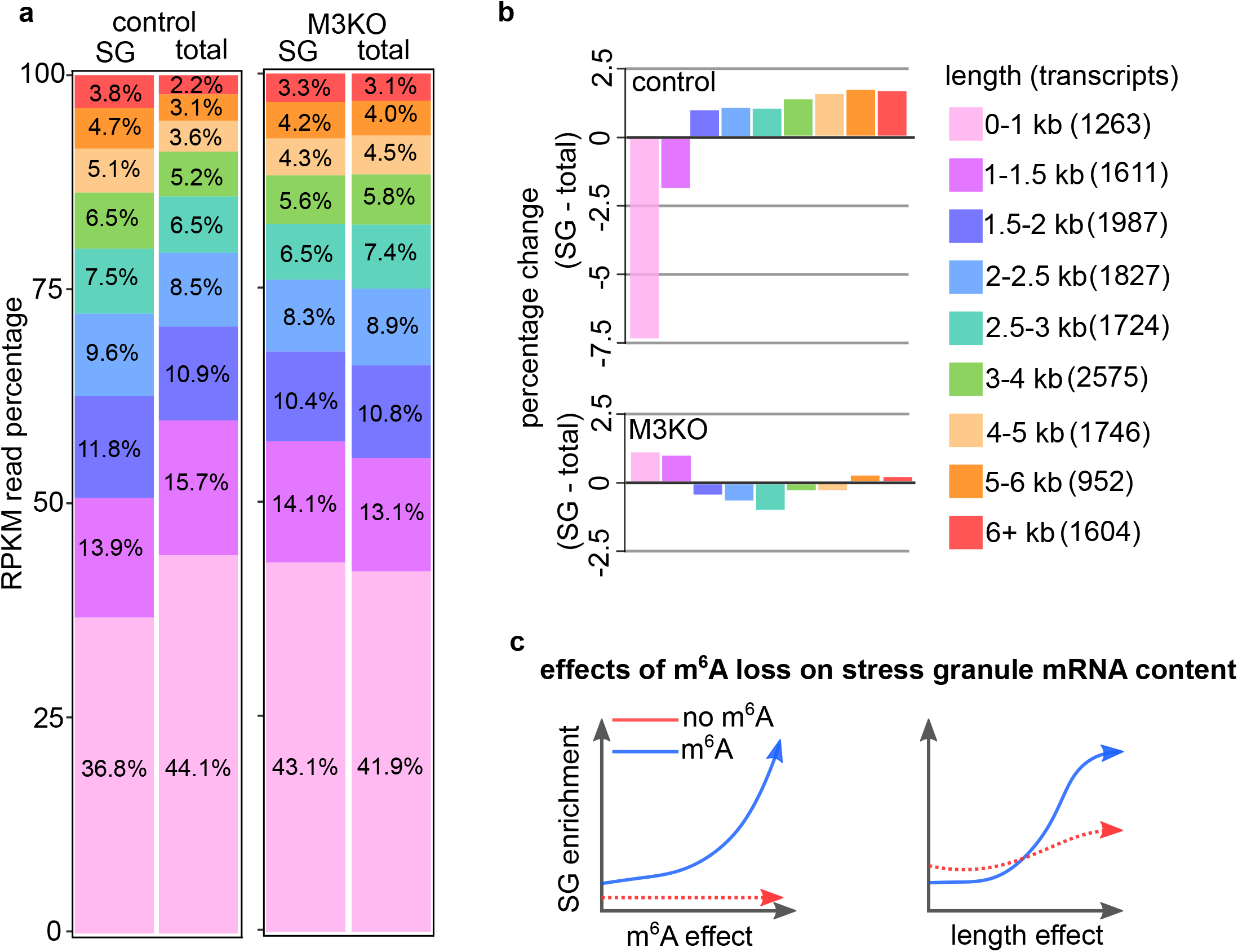
Alternative RPKM comparison of stress granule enrichment of mRNAs based on length. **a,** Comparison of the percentage of reads per kilobase per million (RPKM) between stress granule and total mRNA samples grouped by transcript length. Average RPKM values for mRNAs containing m^6^A sites were calculated and compared across samples. The percentages are approximated to the last digit given. **b,** Percentage change between stress granule and total mRNA RPKM in control and *Mettl3* KO grouped by mapped m^6^A sites. The data shows the percentage differences depicted between each of the categories presented in **a**. The percentages are approximated to the hundredth. In control, a length-dependent enrichment effect is observed for all mRNAs longer than 1.5 kb. In *Mettl3* KO, only mRNAs that are very short (<1.5 kb) or very long (>5 kb) are enriched in stress granules. Notably, shorter mRNAs in the 0-1.5 kb group comprise 50-60% of total overall reads, indicating the abundance of these transcripts. These reads are greatly de-enriched in control stress granules. In contrast, these abundant, short transcripts are enriched in *Mettl3* KO stress granules. **c,** Estimating the effects of mRNA on the stress granule transcriptome. Blue arrowed lines depict observations in control cells, while red arrowed lines represent observations during in *Mettl3* KO. In the left panel, the m^6^A-dependent effect on stress granule enrichment is shown. In cells with METTL3, m^6^A-DF interactions exert a strong effect on mRNA recruitment. The m^6^A effect is absent in cells lacking METTL3. In the right panel, the length effect on stress granule enrichment is shown. Under standard conditions, long RNAs interact with one another through RNA-RNA interactions and multivalent DF-m^6^A effects. In the absence of m^6^A, shorter mRNAs enter stress granules. Many longer mRNAs are affected to a lesser degree by loss of m^6^A owing to compensatory RNA-RNA and RNA-protein interactions.

**Extended Data Fig. 8:**
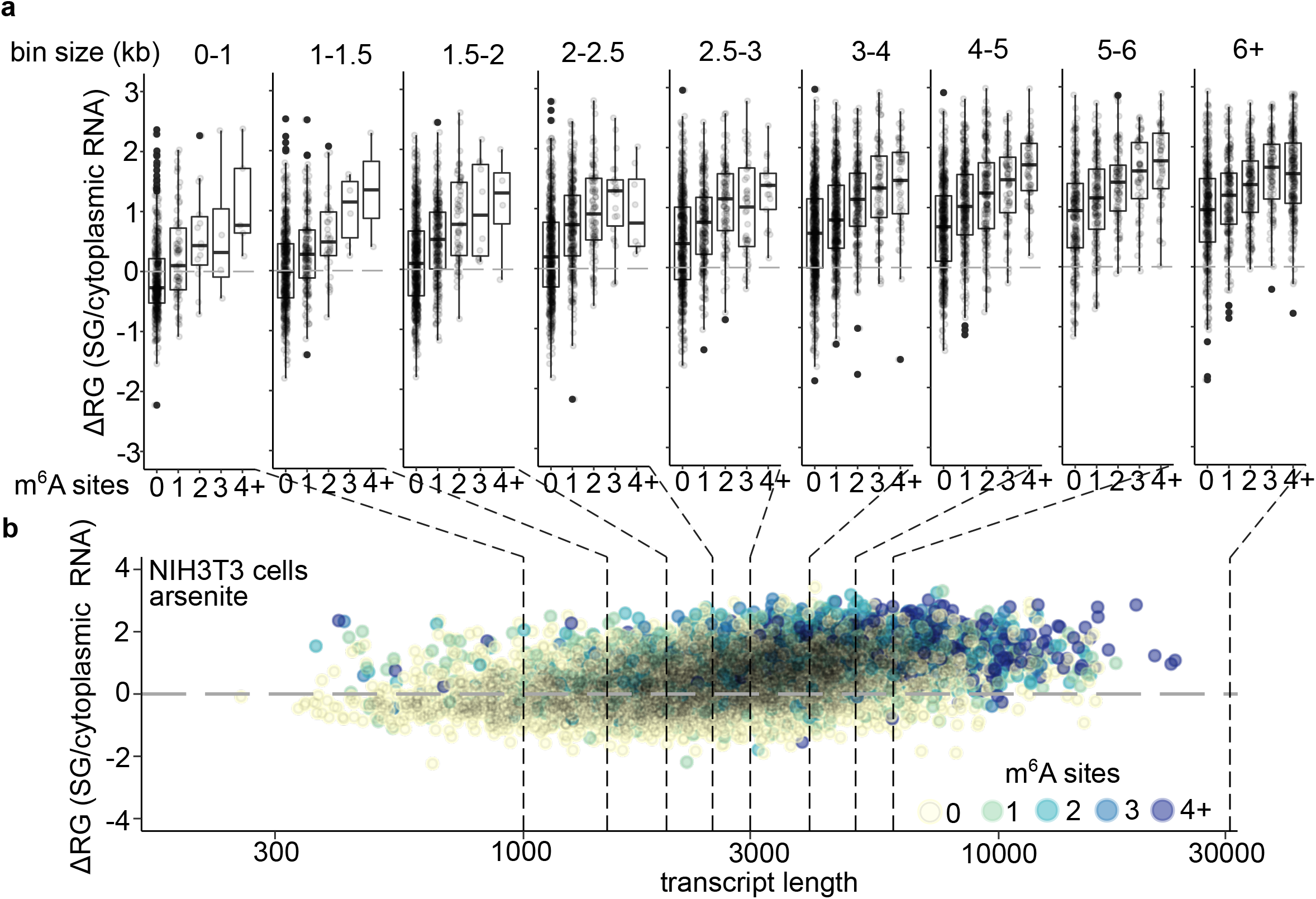
mRNA enrichment as a function of transcript length and m^6^A in NIH3T3 cells. **a,** Boxplots demonstrating the correlation between m^6^A levels and stress granule enrichment in NIH3T3 cells. The y-axis shows the level of enrichment of transcripts in stress granules relative to the cytoplasm. The x-axis shows the number of m^6^A sites per transcript. Boxplots depict the median, upper and lower quartiles. Each dot represents a unique individual transcript. At the top of each boxplot the range of transcript lengths contained in the plot below is shown. Transcript size increases from left to right. Generally, regardless of transcript length, m^6^A has a cumulative effect on the likelihood of enrichment in stress granules. **b,** Scatter plot demonstrating the correlation between m^6^A levels and stress granule enrichment in NIH3T3 cells. The y-axis shows the relative level of enrichment of transcripts in stress granules relative to the cytoplasm. The x-axis shows the length of individual transcripts on a logarithmic scale. The number of m^6^A sites in each individual transcript is color-coded as shown (yellow = 0, light green = 1, blue-green = 2, blue = 3, violet = 4+). Dotted lines are shown connected to the boxplots in panel **a** that summarize the scatter data per m^6^A site.

**Supplementary Table 1**

Differential expression analysis of RNA-seq data from control and *Mettl3* KO cells. Analysis was performed using the DESeq2 package in R^48^. RPKM values provided for each condition are the average of two biological replicates. Short and long transcript isoform lengths were obtained from Ensembl GRCm38.75 (mm10)^46^. m^6^A site counts were obtained from m^6^A-seq data for E13.5 MEFs (GSE61995)^27^.

**Supplementary Table 2**

Alternative polyadenylation (APA) site usage analysis of RNA-seq data from control and *Mettl3* KO cells. Analysis was performed using the DaPars algorithm in Python^40^. Raw RNA-seq counts from two biological replicates were used as input.

**Supplementary Table 3**

The m^6^A-dependent fold change enrichment (ΔFCE) of transcripts in *Mettl3*- and m^6^A-depleted cells relative to control cells. A positive ΔFCE indicates lower dependence on m^6^A for stress granule enrichment, while a negative ΔFCE indicates higher dependence on m^6^A for stress granule enrichment.

